# A lncRNA-mediated metabolic rewiring of cell senescence

**DOI:** 10.1101/2024.01.25.577153

**Authors:** Elena Grossi, Francesco P. Marchese, Jovanna González, Enrique Goñi, José Miguel Fernandez-Justel, Alicia Amadoz, Nicolás Herranz, Leonor Puchades-Carrasco, Marta Montes, Maite Huarte

## Abstract

Despite the classical view of senescence as passive growth arrest, senescent cells remain metabolically active to be able to cope with the energetic demand of the senescence program. However, the mechanisms underlying this metabolic reprogramming remain poorly understood. We have identified *sin-lncRNA,* a previously uncharacterized lncRNA, that plays a pivotal role in this response. *Sin-lncRNA* is only expressed by senescent cells, induced by the senescence master regulator C/EBPβ. While strongly activated in senescence, *sin-lncRNA* loss reinforces the senescence program by altering oxidative phosphorylation and rewiring mitochondrial metabolism. By interacting with the TCA enzyme dihydrolipoamide S-succinyltransferase (DLST) it facilitates its localization to the mitochondria. On the other hand, s*in-lncRNA* depletion results in DLST nuclear translocation linked to DLST-dependent transcriptional alteration of OXPHOS genes. While in highly proliferative cancer cells, *sin-lncRNA* expression remains undetected, it is strongly induced upon cisplatin-induced senescence. Depletion of *sin-lncRNA* in ovarian cancer cells results in deficient oxygen consumption and increased extracellular acidification, sensitizing the cells to cisplatin treatment. Altogether, these results indicate that *sin-lncRNA* is specifically induced in cellular senescence to maintain metabolic homeostasis. Our findings reveal a new regulatory mechanism in which a lncRNA contributes to the adaptive metabolic changes in senescent cells, unveiling the existence of an RNA-dependent metabolic rewiring specific to senescent cells.

## INTRODUCTION

Senescence is a permanent state of growth arrest that can be induced by different types of stress, such as shortening of telomeres due to extensive replication, DNA damage, oxidative stress, chemotherapy treatment or oncogene overexpression ^1^. Senescent cells present characteristic morphological and biochemical features such as enlarged size, halted proliferation, activation of senescence-associated (SA) β-galactosidase activity and increased expression of cell cycle inhibitors ^1^. Furthermore, in several types of senescence the cells secrete a combination of interleukins, metalloproteases and growth factors, collectively known as the senescence-associated secretory phenotype (SASP) ^2–4^. While the SASP reinforces senescence, its chronic activation can have pro-tumorigenic effects and contribute to several aging related pathologies ^5^. In fact, the persistence of therapy-induced senescent cancer cells in the tumor has been linked to resistance to chemotherapy ^6, 7^. There is, therefore, a critical need to develop combinatorial approaches to not only stop proliferation, but also induce the elimination of the senescent tumor cells.

Despite their decreased proliferative potential, senescent cells are metabolically active in order to cope with the energetic demands of the senescence program ^8^. Indeed, metabolic reprogramming is considered a hallmark of senescence ^9^. In general, senescent cells accumulate dysfunctional mitochondria resulting in an increase of reactive oxygen species (ROS)^10–12^. Although different triggers of senescence result in broad metabolic differences, the interplay between senescent cells and metabolism is highly dynamic and context dependent, and the underlying mechanisms remain largely unexplored. In this context, long noncoding RNAs (lncRNAs) represent a relatively uncharted territory for investigation.

LncRNAs are transcripts recently defined as longer than 500 nucleotides with exquisite cell-type specific expression patterns that lack protein-coding potential ^13^. They are tightly regulated during development or in response to signaling pathways ^14^. Although it is still unclear how many of the thousands of annotated lncRNAs have a significant biological role, several have been found to be essential for the regulation of key cellular processes, such as proliferation and differentiation, as well as a broad range of diseases including cancer ^15, 16^. Notably, our previous work and other laboratories’ have reported that specific lncRNAs are differentially expressed during senescence induction ^17, 18^, regulating multiple senescence aspects, from transcriptional response to SASP production ^19–22^. Intriguingly, recent studies have involved a number of lncRNAs in the regulation of cellular metabolism ^23–26^, and ncRNAs are emerging as interactors and regulators of metabolic enzymes ^27^. However, the contribution of this functional diverse class of molecules to the metabolic reprogramming of senescent cells remains unknown.

Here we report the identification of the senescence-specific lncRNA, *sin-lncRNA*, which is induced in senescence to prevent uncontrolled metabolic alterations, by regulating the function of a key metabolic enzyme and the transcription of metabolic genes. Together, our results provide evidence of an RNA-dependent metabolic rewiring specific to the senescent cellular state.

## RESULTS

### Oncogene-induced senescence leads to broad changes in gene expression, encompassing alterations in lncRNAs

Although the importance of lncRNAs in a variety of cellular processes is now fully recognized, only a handful of them have been described as directly implicated in cellular senescence. To identify novel regulators of this process, we analyzed RNA-seq data from a cellular system that recapitulates senescence induction *in vitro* thanks to the controlled activation of an oncogene ^28^. IMR90 non-immortalized lung fibroblasts were engineered to express an oncogenic HRAS variant (HRAS^G12V^) fused to the estrogen receptor (ER:RAS), which is activated following 4-hydroxy-tamoxifen (4OHT) administration (Figure 1A) ^3, 29^. Multiple senescence markers, such as arrest of proliferation, increased expression of pro-inflammatory factors and tumor suppressors along with detectable β-galactosidase staining, were observed, ensuring a proper senescence progression (Suppl. Figure 1A-E). Furthermore, the presence of senescence-associated heterochromatin foci (SAHF) as well as an increase in H3K9me3 and γH2AX foci, confirmed the chromatin remodeling linked to oncogene-induced senescence (OIS) (Suppl. Figure 1F, G). RNAseq data generated in this cell system ^28^, detected more than 4720 genes differentially expressed upon induction of senescence (log2FC>1; padj<0.05), including not only many coding genes (3456) but also lncRNAs (951) (Figure 1B and Suppl. Figure 1H). The majority of lncRNAs (62%) were downregulated, which might reflect cell cycle inhibition during senescence, whereas the remaining 38% of lncRNAs were upregulated. We focused our attention on lncRNAs highly induced upon OIS, as we speculated, they might play a role in the senescence process, and we verified their upregulation at different time points of OIS progression (Figure 1C and Suppl. Figure 1I).

**Figure 1.**
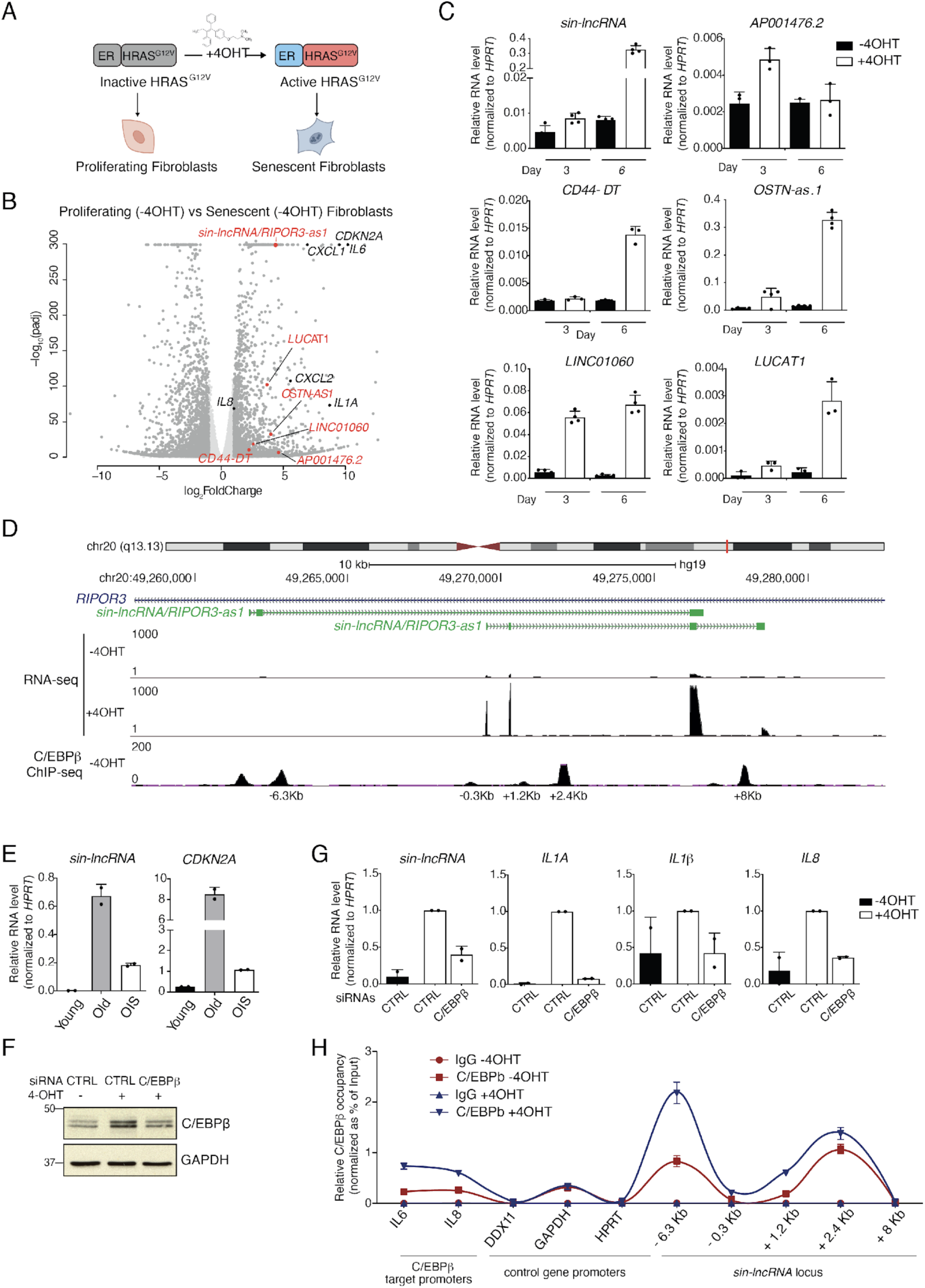
*sin-lncRNA* is a senescence-specific lncRNA transcriptionally induced by C/EBPβ. **A)** Schematic representation of normal fibroblasts undergoing OIS upon 4OHT-mediated activation of an oncogenic form of HRAS. **B)** Volcano plot depicting differential gene expression data (GSE61130) comparing IMR90 ER:RAS proliferating and senescent cells. Cut-off for significance log_2_FC>±1; adj.Pvalue < 0.05. A selection of canonical senescence markers (in black) and lncRNAs (in red) is shown. **C)** RT-qPCR analysis of a set of lncRNA candidates upregulated upon 4OHT time course (3 or 6 days) in IMR90 ER:RAS cells; *RIPOR3-as1* was amplified with primer set#2 (see Suppl. Figure 2A for details) Data are presented as mean values ± s.d. n=3/4. **D)** Genomic snapshot of *sin-lncRNA* locus with RNA-seq tracks of untreated and 4OHT-treated IMR90 ER:RAS fibroblasts, as well as C/EBPβ ChIP-seq ENCODE data generated in IMR90 cells. **E)** RT-qPCR analysis of “young” (<14 passages), “old” (>30 passages) IMR90 fibroblasts and 4OHT-induced “OIS” IMR90 ER:RAS cells, shows that *sin-lncRNA* upregulation is maintained in replicative senescence, compared to p16 positive control. (n=2) **F)** Western blot analysis of control and C/EBPβ-depleted senescent IMR90 ER:RAS cells. Proliferating fibroblasts were used as further control. **G)** RT-qPCR analysis of *sin-lncRNA*, as well as a set of C/EBPβ-target genes in control and C/EBPβ-depleted senescent IMR90 ER:RAS cells. Data are represented as mean values ± s.d. (n=2) **H)** ChIP-qPCR analysis of C/EBPβ in proliferating and senescent IMR90 ER:RAS cells upon 6 days of 4OHT treatment. IgG IP was used as IP control; C/EBPβ peaks at *sin-lncRNA* locus are labeled according to their distance from *sin-lncRNA* TSS; regions located in *DDX11*, *GAPDH* and *HPRT* promoters served as negative controls, *IL6* and *IL8* promoter regions as positive controls (n=2).

Interestingly, we observed a previously uncharacterized lncRNA (*ENSG00000234693)* as one of the most significantly and strongly upregulated genes, with an induction comparable to that of key senescence genes *CKN2A, CXCL1* and *IL6* (Figure 1B and Suppl. Table 1). We consequently renamed it *senescence-induced lncRNA* or *sin-lncRNA*. *sin-lncRNA* is an intronic lncRNA located in chromosome 20 and transcribed in the antisense orientation of the protein-coding gene *RIPOR3* (Figure 1D, Suppl. Figure 2A, B). *sin-lncRNA* lacks coding potential (Suppl. Figure 2C) and presents two poly-adenylated isoforms annotated by GENCODE (v41); a longer one (638 nts), composed by three exons, and a shorter one (539 nts) of four exons. However, only the shorter isoform is expressed and highly upregulated upon 4OHT treatment in IMR90 fibroblasts, according to both sequencing data and RT-qPCR performed with different primer sets (Figure 1D and Suppl. Figure 2A). This observation was confirmed by absolute quantification by RT-qPCR, which showed that *sin-lncRNA* is lowly expressed in proliferating conditions with ∼9 copies per cell, while, upon senescence induction, the expression of its short isoform increases up to ∼86 copies (Suppl. Figure 2D).

Interestingly, *sin-lncRNA* levels increased only at later stages of senescence (Figure 1C and Suppl. Figure 2A), prompting us to investigate whether its upregulation was dependent on the full activation of a senescence program or rather related to the induction of the HRAS oncogene in OIS. To this aim, we analyzed *sin-lncRNA* expression levels in different types of senescence, such as replicative senescence – achieved by prolonged culture of untransformed IMR90 fibroblasts – and senescence induced by ionizing radiation (γ-irradiation). In all the systems analyzed *sin-lncRNA* expression was strongly induced in the senescent context, correlating with known senescence markers, such as *CDKN2A* and *IL6* (Figure 1E and Suppl. Figure 2E).

Due to the strong and highly specific *sin-lncRNA* induction in senescence, we speculated that senescence-specific factors could regulate the expression of this lncRNA. We took advantage of ChIP-seq data generated in IMR90 ^30^ cells to examine *sin-lncRNA* genomic region finding that the transcription factor C/EBPβ was bound to several positions in the proximity of *sin-lncRNA* transcription start site (TSS) and gene body (Figure 1D). Interestingly, C/EBPβ exerts an important role in the later stages of senescence induction, mainly by activating genes related to inflammation and SASP ^31^. After confirming the previously reported C/EBPβ upregulation in OIS ^2^ (Suppl. Figure 2F), we depleted C/EBPβ in senescent cells and analyzed *sin-lncRNA* expression (Figure 1F, G and Suppl. Figure 2G). Notably, the induction of *sin-lncRNA* in senescence was prevented by C/EBPβ depletion, similarly to what occurred to *IL1β*, *IL6* and *IL8,* other canonical targets of this transcription factor (Figure 1G). To further investigate whether or not *sin-lncRNA* is a C/EBPβ transcriptional target, we performed ChIP-qPCR of this transcription factor in proliferative and senescent conditions (Figure 1H). C/EBPβ was found enriched at *sin-lncRNA* regulatory regions and its binding increased in senescent cells (Figure 1H), supporting the evidence that the senescence driver C/EBPβ controls *sin-lncRNA* expression at later stages of senescence induction and suggesting that the lncRNA might exert a role in this response.

### *sin-lncRNA* loss contributes to the senescent phenotype through increase of oxidative stress

To explore the function of *sin-lncRNA*, we depleted it in senescent IMR90 ER:RAS fibroblasts (Suppl. Figure 3A) and analyzed several senescence markers at 5 days of induction to detect intermediate states before the cells enter a fully stable senescent phenotype ^32^. Interestingly, *sin-lncRNA* KD further decreased proliferation of senescent cells (Figure 2A, C and Suppl. Figure 3B), increased the percentage of β-galactosidase positive cells (Figure 2C) and activated the p53 signaling pathway (Figure 2D), without changing levels of apoptosis (Suppl. Figure 3C), indicating that *sin-lncRN*A knockdown reinforces the senescent phenotype.

**Figure 2.**
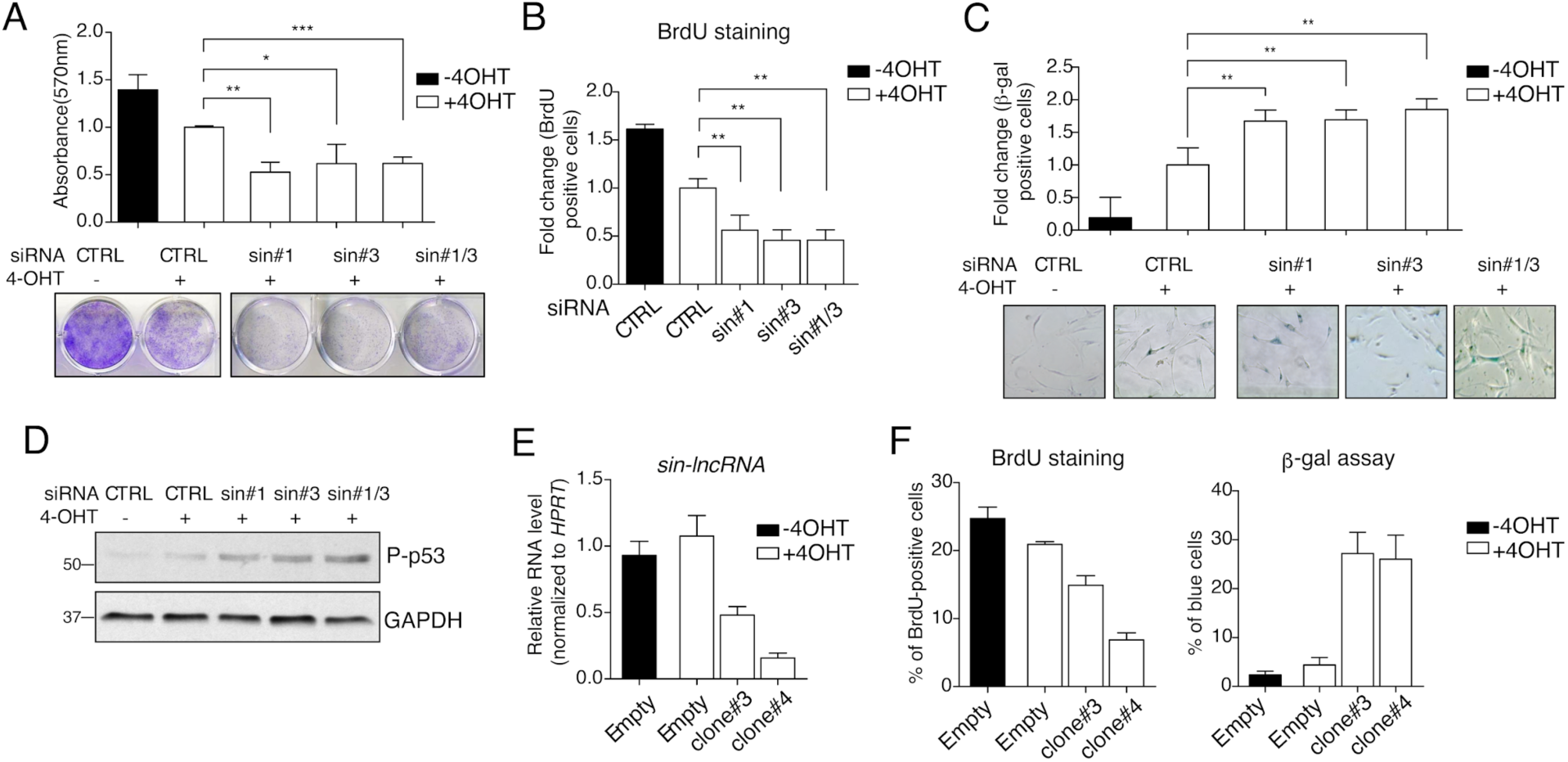
*sin-lncRNA* depletion reinforces the senescent phenotype. **A)** Crystal violet assay and **B)** BrdU analysis assessing cell growth of control and *sin-lncRNA* depleted IMR90 ER:RAS fibroblasts after 5 days of senescence induction. **C)** β-galactosidase analysis of control and *sin-lncRNA* depleted IMR90 cells induced to senescence for 8 days. **D)** Western blot analysis of control and *sin-lncRNA* depleted IMR90 cells treated with 4OHT for 5 days. **E)** RT-qPCR analysis of TIG3 ΔBRAF cell clones infected with *sin-lncRNA* sgRNA or an empty plasmid. **F)** BrdU (left) and β-galactosidase (right) assays of control or *sin-lncRNA* sgRNA TIG3 ΔBRAFclones.

To investigate the broader significance of these results, we investigated other senescence systems and methods for *sin-lncRNA* loss-of-function. We took advantage of TIG3 hTERT lung fibroblasts engineered to express an inducible and constitutively active portion of the human BRAF oncogene to drive OIS (TIG3 ΔBRAF:ER) ^33^. In this cell system, we again observed reduced proliferation upon *sin-lncRNA* downregulation by RNAi (Suppl. Figure 3D-F), indicating that *sin-lncRNA* depletion leads to the same phenotype regardless of the type of oncogene expressed. We also generated TIG3 ΔBRAF:ER cell lines stably depleted of *sin-lncRNA* by CRISPR/Cas9 technology. Despite not being able to isolate viable cell clones with *sin-lncRNA* biallelic deletion, we found that the removal of just one allele was sufficient to decrease *sin-lncRNA* expression, leading to a consistent strengthening of the senescence phenotype (Figure 2E-F and Suppl. Figure 3G-H). Altogether, these results suggest that the loss of *sin-lncRNA* reinforces the induction of cellular senescence.

To investigate which aspect of the complex senescence response is influenced by *sin-lncRNA*, we analyzed the global changes in gene expression by RNA-seq of *sin-lncRNA*-depleted ER:RAS fibroblasts upon senescence induction (Figure 3A, Suppl. Figure 4A and Suppl. Table 2), and employed Gene Set Enrichment Analysis (GSEA) to determine enriched pathways associated with the observed changes in gene expression (Figure 3B and Suppl. Figure 4B). Interestingly, among the most significantly reduced terms were several metabolic pathways, such as cholesterol homeostasis and fatty acid metabolism. Moreover, mRNAs belonging to oxidative phosphorylation pathway were significantly downregulated, including the mitochondrial ADP/ATP carrier ACC 1, 2, 3 (*SLC25A4*, *SLC25A5*, *SLC25A6*, respectively), which regulate key transport steps for cellular ATP production ^34^ (Figure 3B-C).

**Figure 3.**
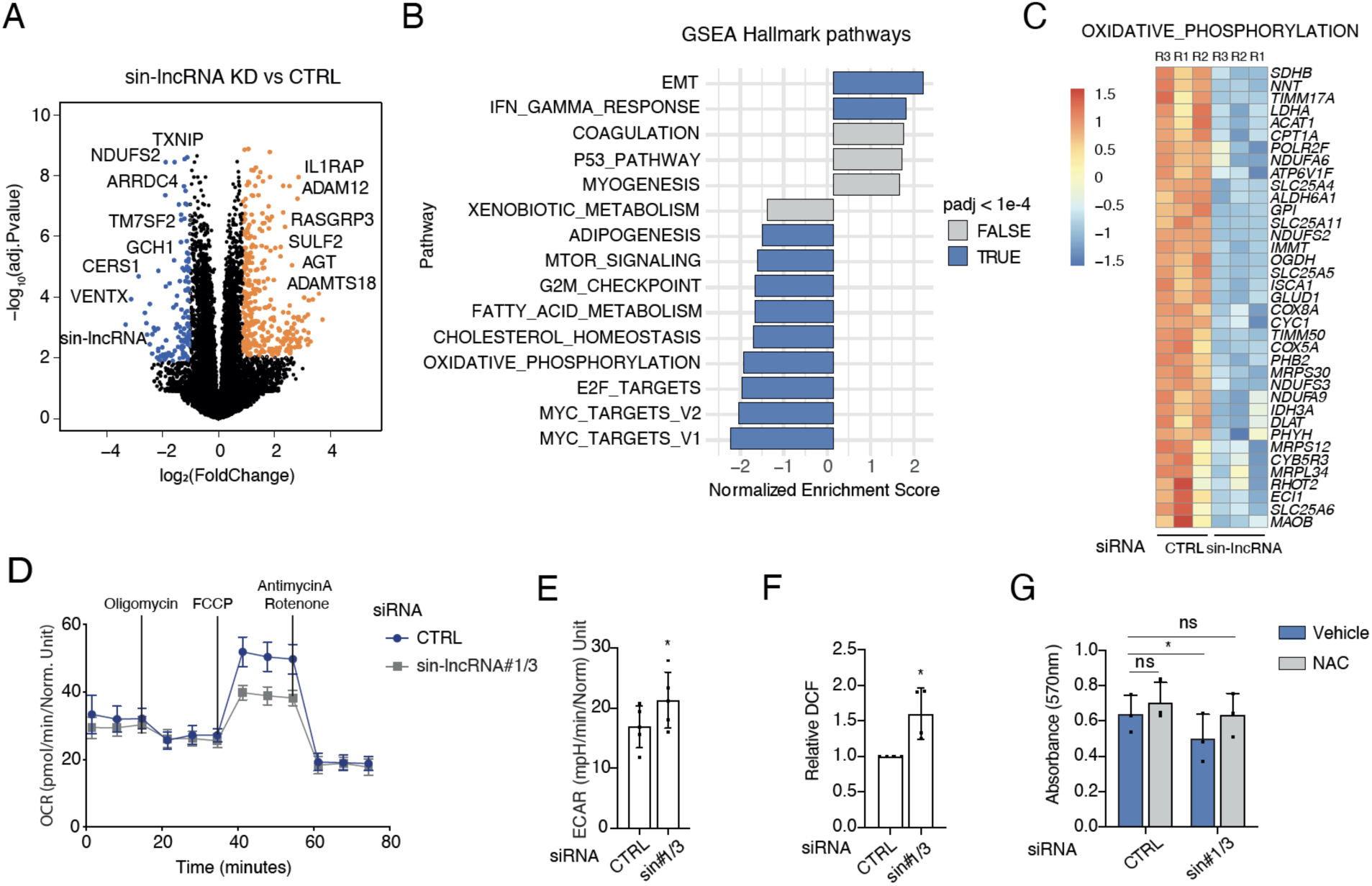
*sin-lncRNA* KD alters oxidative stress response and metabolic pathways in senescent cells. **A)** Volcano plot depicting differentially expressed genes comparing *sin-lncRNA* KD vs control IMR90 ER:RAS senescent cells. Upregulated genes (adj.Pvalue < 0.01; log2FC>1) are labeled in red, downregulated genes (adj.Pvalue < 0.01; log2FC<-1) in blue. **B)** Hallmark Gene Set Enrichment Analysis (GSEA) reporting the most enriched pathways associated to the genes differentially expressed (DE) upon *sin-lncRNA* depletion (threshold: adj.Pvalue<1e-4). **C)** Heatmap highlighting expression changes across RNA-seq replicates of genes belonging to OXIDATIVE_PHOSPHORILATION GSEA pathway. **D)** Oxygen Consumption Rate (OCR) of control or *sin-lncRNA* depleted IMR90 ER:RAS senescent cells obtained using the Seahorse XFe96 Analyzer following injections of oligomycin, FCCP, and rotenone/antimycin A, as indicated. Data is presented as mean ± s.d. (n = 4). 2-way ANOVA and Sidak’s multiple comparisons test, *\*p<*0.05. **E)** Basal extracellular acidification rate (ECAR) measured using the Seahorse XFe96 Analyzer, in the same conditions as in D, and following injections of glucose, oligomycin, and 2-deoxy-D-glucose, as indicated in the Glycolysis Stress Test. Data is presented as mean ± s.d. (n = 4). Two tailed student’s *t*-test, *\*p<*0.05. **F)** Fluorometric analysis measuring the amount of the fluorescent oxidized probe 2′,7′-dichlorofluorescein (DCF), as indicative of ROS/NOS intracellular levels in control and *sin-lncRNA* depleted IMR90 ER:RAS fibroblasts induced to senescence for 5 days. Proliferating control cells were used as further control. **G)** Absorbance levels of crystal violet solution measured at 560nm from vehicle or N-acetyl-L-cisteine (NAC)-treated IMR90 ER:RAS fibroblasts transfected with siRNA control (CTRL) or against *sin-lncRNA* (sin#1/3) and induced to senescence for 5 days. The graph represents the mean ± s.d. (n=3). Two tailed student’s *t*-test, *\*p<*0.05.

Since metabolic rearrangements and altered oxidative phosphorylation (OXPHOS) are known to be linked to senescence ^13, 35^, we speculated that *sin-lncRNA* could influence the metabolic states of senescent cells. To analyze metabolic alterations upon *sin-lncRNA* depletion, we measured the oxygen consumption rate (OCR) coupled to ATP production during OXPHOS by Seahorse assay ^36^. Compared to control senescent cells, *sin-lncRNA* knockdown led to the reduction of several key parameters of mitochondrial function, most significantly impacting maximal respiration rate, mitochondrial spare capacity and coupling efficiency (Figure 3D, and Suppl. Figure 4C). We also observed an increment in extracellular acidification rate (ECAR) in *sin-lncRNA depleted cells*, a reflection of increased aerobic glycolysis that has been linked to defects in ATP production ^37^ (Figure 3E).

Since alterations in OXPHOS are also linked to increased oxidative stress ^38–40^, we measured intracellular levels of reactive oxygen and nitrogen species (ROS/RNS) and observed a significant enrichment of oxidant species in *sin-lncRNA*-depleted senescent ER:RAS cells (Figure 3F), suggesting that *sin-lncRNA* is involved in the control of oxidative stress in senescent cells. The increase in oxidant species and metabolic dysfunctions can reinforce senescence ^41, 42^. We therefore hypothesized that *sin-lncRNA* depletion might exacerbate senescence-related mitochondrial alteration and promote oxidative stress, thereby leading to the observed senescence reinforcement. Indeed, blocking ROS accumulation by N-acetyl-L-cysteine (NAC) treatment ^43, 44^ restored cell growth and reduced SA-β-galactosidase activation in senescent cells depleted for *sin-lncRNA* (Figure 3G and Suppl. Figure 4D). Together these data support the notion that *sin-lncRNA* contributes to manage the metabolic rewiring and oxidative stress response of senescent cells.

### *sin-lncRNA* interacts with DLST and controls its subcellular localization

LncRNAs have been previously associated with defects in oxidative stress and metabolic pathways, in particular in cancer, where they alter gene programs through different mechanisms ^19, 45^. To elucidate how *sin-lncRNA* regulates metabolic response of senescent cells, we first analyzed *sin-lncRNA* subcellular localization by RNA-FISH and found it to be mainly enriched in the cytoplasm of senescent fibroblasts (Figure 4A and Suppl. Figure 5A). Interestingly, an independent study ^46^ indicated that *sin-lncRNA* is enriched in the mitoplasts of senescent cells (Suppl. Figure 5B). We purified mitochondria from senescent cells by FACS sorting, and detected the presence of *sin-lncRNA* along with mitochondrially encoded transcripts (Figure 4B and Suppl. Figure 5C). Together, these data suggest that *sin-lncRNA* acts in the mitochondria to control proliferative and metabolic pathways of senescent cells.

**Figure 4.**
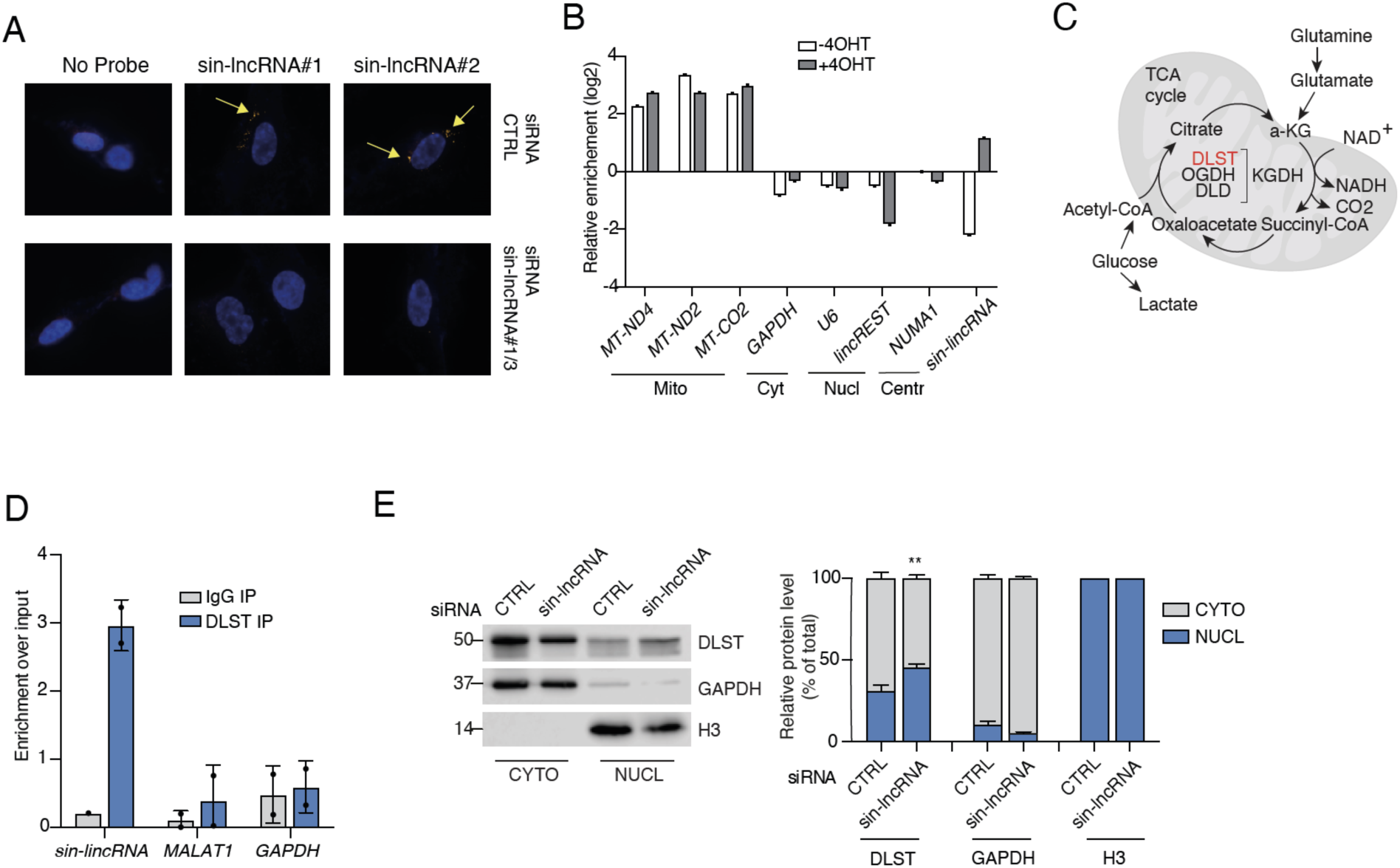
*sin-lncRNA* interacts with DLST and controls its subcellular localization. **A)** RNA-FISH based on locked nucleic acid (LNA) technology on fixed IMR90 ER:RAS cells depleted or not of *sin-lncRNA* (siRNA_sin-lncRNA#1/3) and treated for 5 days with 4OHT. Two different LNAs were used. Incubation without LNA probe (“No_probe”) was used as control. **B)** RT-qPCR analysis of a set of genes after mitochondria purification from proliferating (–4OHT) or senescent cells (+4OHT) using Mitotracker (green, 488) staining and FACS sorter. The graph shows a representative experiment. **C)** Schematic representation of the TCA cycle that takes place in the mitochondria. DLST enzyme is highlighted in red. **D)** RNA immunoprecipitation analysis monitoring DLST binding to *sin-lncRNA, MALAT1* and *GAPDH* RNA in control and *sin-lncRNA* depleted (siRNA#1/3) senescent IMR90 ER:RAS, using anti-DLST antibody or IgG as a negative control. The graph represents the mean ± s.d. (n=2). **E)** *Left*, western blot analysis of DLST cellular fractionation in control and *sin-lncRNA* depleted (siRNA#1/3) senescent IMR90 ER:RAS. GAPDH and H3 were used as cytoplasmic and nuclear controls respectively. *Right*, percentage of nuclear and cytoplasmic distribution of DLST normalized to GAPDH (cyto) or H3 (nucl). The graph represents the mean ± s.d. (n=3). Two tailed student’s *t*-test, \**\*p<*0.01.

Reasoning that *sin-lncRNA* function could be mediated through interacting proteins, we set to unbiasedly identify its protein partners by RNA pulldown. We incubated *in vitro* transcribed *sin-lncRNA* or an unrelated control RNA with senescent IMR90 ER:RAS extracts and isolated interacting proteins (material and methods) (Suppl. Figure 5D). Mass spectrometry (MS) revealed several proteins enriched (>2 unique peptides) specifically associated with *sin-lncRNA* but not to the control RNA (Suppl. Figure 5E and Suppl. Table 3). Notably, the most abundant proteins retrieved by MS analysis were Dihydrolipoamide S-succinyltransferase (DLST) and DEAD-Box Helicase 28 (DDX28), which localize in the mitochondria and have a recognized role in that compartment ^47^, in line with our findings linking *sin-lncRNA* to senescent cell metabolism.

DLST, identified as the main *sin-lncRNA* interactor, is the E2 component of the α-ketoglutarate (αKG) dehydrogenase complex (KGDH), which is essential for the entry of glutamine into the tricarboxylic acid (TCA) cycle ^48^ (Figure 4C). DLST has been recently identified as an RNA binding protein in an RNA binding screen ^49^. We orthogonally confirmed the interaction between the endogenous *sin-lncRNA* and DLST by RNA Immunoprecipitation in senescent fibroblasts (Figure 4D and Suppl. Fig 5F), suggesting they are functionally related. In order to map the region of *sin-lncRNA* interacting with DLST, we performed *in vitro* pulldown with biotinylated fragments spanning the 539 nucleotides of the full-length transcript. We observed that all fragments except F1 (1-150 nts) are involved in the binding (Suppl. Fig 5G).

We then sought to investigate whether *sin-lncRNA* would regulate DLST expression or stability. However, DLST mRNA and protein levels were not affected upon *sin-lncRNA* knockdown and remained stable upon senescence induction (Suppl. Figure 5H, I), indicating that neither *sin-lncRNA* nor cellular senescence modulate DLST expression. Of note, DLST is a protein that shuttles between different subcellular compartments. While its main activity takes place in the mitochondria, it is also localized in the nucleus, where it is known to regulate gene expression ^50–52^. We therefore investigated whether DLST localization might be regulated by the lncRNA. Interestingly, cellular fractionation of senescent cells upon *sin-lncRNA* depletion showed that DLST levels were significantly decreased in the cytoplasm and increased in the nuclear fraction of cells (Figure 4E) in contrast to control senescence conditions (Suppl. Figure 5J), suggesting that *sin-lncRNA* controls the subcellular localization of the metabolic enzyme DLST.

### *sin-lncRNA* rewires metabolic activity in senescent cells

Given the change of DLST localization observed upon *sin-lncRNA* depletion, we sought to compare the phenotype of DLST depletion with the phenotypic alterations observed upon *sin-lncRNA* knockdown in senescent cells, as previous reports demonstrated increased ROS production upon DLST knockdown ^53^. DLST knockdown in senescent fibroblasts led to an increase in the percentage of β-galactosidase positive cells and decreased cell growth (Figure 5A, B), thus phenocopying in this regard *sin-lncRNA* knockdown. Deeper analysis of metabolic alterations showed an increase in lactate secretion in both *sin-lncRNA* and DLST-depleted senescent cells compared to control cells (Figure 5C), probably as a result of higher glycolytic activity in these cells (Figure 3C).

**Figure 5.**
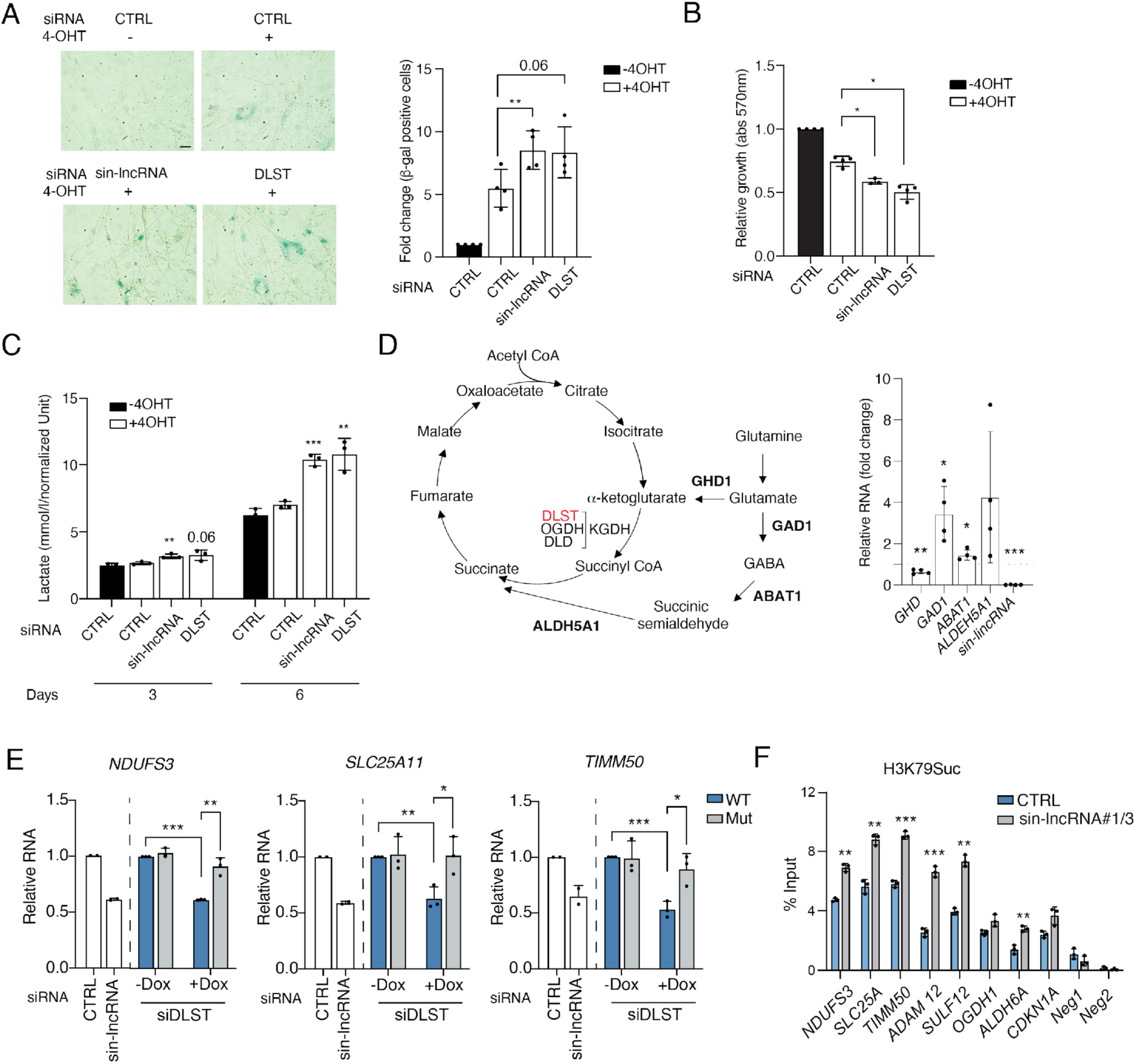
*sin-lncRNA* rewires metabolic activity in senescent cells. **A)** *Left*, representative β-galactosidase images of untreated control cells and control, *sin-lncRNA* depleted or DLST-depleted IMR90 cells induced to senescence for 5 days. Scale bar: 50 μm. *Right*, quantification of the percentage of β-galactosidase positive cells in each condition normalized to control untreated cells set as 1. The graph represents the mean ± s.d. (n=4). Two tailed student’s *t*-test, *\*\*p<*0.01. **B)** Absorbance levels of crystal violet solution measured at 560nm in the conditions described in E. The graph represents the mean ± s.d. (n=4). Two tailed student’s *t*-test, *\*p<*0.05. **C)** Lactate levels measured from the supernatant of IMR90 ER:RAS fibroblasts transfected with the indicated siRNA and induced with 4OHT for 3 or 6 days. The graph represents the mean ± s.d. (n=3). Two tailed student’s *t*-test, *\*p<*0.05; **p<0.01; ***p<0.001. **D)** Left panel, schematic showing alternative metabolic pathways. Right panner, qRT-PCR of the corresponding transcripts in sin-lncRNA knockdown cells relative to control. (n=3). The graph represents the mean ± s.d. (n=4). Two tailed student’s *t*-test, *\*p<*0.05; **p<0.01; ***p<0.001. **E)** qRT-PCR of metabolic transcripts in the cells in stable senescent IMR90 ER:RAS cells expressing a doxycycline-inducible wild type (WT) or NLS mutant (Mut) (R224A/K226E) DLST construct in the absence (-Dox) or presence (+Dox) of 100ng/ml of doxycycline for 72h (n=3). **F)** ChIP-qPCR analysis of H3K79 Succinylation in senescent IMR90 ER:RAS cells upon 6 days of 4OHT treatment transfected with siRNA control or siRNA against *sin-lncRNA* at the promoters of different metabolic genes. Regions located at intergenic regions (Neg1 and Neg2) served as negative controls. The graph shows a representative experiment (n=2).

We further investigated the role of *sin-lncRNA* in metabolism by comparing changes in the levels of glutamine-derived metabolites upon *sin-lncRNA* and DLST knockdown in senescent cells using Liquid Chromatography – Mass Spectrometry (LC-MS) and fully labeled [U-^13^C] glutamine as a tracer (Figure 5D). The glutamine labeled fraction in cells grown with [U-^13^C] glutamine consisted of >90% M+5 glutamine in all conditions, demonstrating equal incorporation of the labeled substrate (Suppl. Figure 6A). While DLST knockdown strongly affected the TCA cycle, by decreasing the rate of glutamine oxidative metabolism, depletion of *sin-lncRNA* did not alter either oxidative or reductive TCA cycle activity (Suppl. Figure 6A, B and Suppl. Table 4). These results indicate that depletion of *sin-lncRNA* does not fully reproduce the metabolic effects of the knockdown of DLST, as expected, based on the protein relocalization caused by *sin-lncRNA* downregulation.

We hypothesized that, upon *sin-lncRNA* knockdown, alternative mechanisms might intervene to maintain a metabolic balance and compensate for deficiencies in KGDH complex activity. In agreement with this idea, the expression levels of the main enzymes implicated in the GABA shunt, an alternative route for the conversion of glutamate to succinate bypassing the TCA cycle ^54^ were upregulated in *sin-lncRNA* knockdown cells (Figure 5D). We thus reasoned that the lack of the lncRNA resulted in a phenotypic output consequence of the decrease of DLST mitochondrial levels combined with events driven by increased nuclear levels of the protein.

DLST harbors a nuclear localization signal (NLS) that allows its translocation to the nucleus promoting the succinylation of H3K79 ^52^. Of note, we observed a significant overlap between mRNAs regulated by *sin-lncRNA* and genes regulated by KGDH-mediated H3K79 succinylation according to a prior study ^52^ (Suppl. Figure 6C). We therefore investigated if DLST translocation could be involved in the observed transcriptional changes. Interestingly, overexpression of the wild type DLST in senescent cells decreased the transcription of a subset of metabolic mRNAs, mimicking the effect of the depletion of *sin-lncRNA* (Fig 5E and Suppl. Figure 6E, F). In contrast, a mutant DLST that is not able to enter the nucleus, (R224A/K226E) ^52^ had no effect (Figure 5E). In fact, a previous study showed that relevant metabolic genes present H3K79Succ chromatin mark^52^ (Suppl. Figure 6D). Strikingly, we observed that H3K79Succ increased on these genes upon *sin-lncRNA* depletion in senescent cells, correlating with an increase in nuclear DLST (Figure 5F). These results show that the nuclear, but not the cytoplasmic form of DLST, was able to reproduce the gene expression changes observed upon depletion of *sin-lncRNA*, supporting that the nuclear shuttling of DLST causes compensatory transcriptional changes observed in *sin-lncRNA* knockdown condition.

### *sin-lncRNA* regulates OXPHOS in ovarian cancer cells

Our results indicate that *sin-lncRNA* is a regulator of the metabolic state of the senescent cells. To gain insight into the potential significance of *sin-lncRNA* function in a physio-pathological context, we interrogated gene expression data of tumor samples available through The Cancer Genome Atlas (TCGA). *sin-lncRNA* presented low expression across cancer types (Suppl. Figure 7A.), its expression negatively correlated with that of genes involved in active cell division, such as cyclins, as *CCNA2*, and members of the origin replication complex, such as *ORC1* and *MCM6* ^55^ (Suppl. Figure 7B). Accordingly, terms related to cell cycle and mitosis were the most enriched Gene Ontology pathways associated with these genes (Suppl. Figure 7C). Since all these cell division-related genes presented a negative correlation with the senescence-specific *sin-lncRNA*, we speculated that a high expression of this noncoding transcript would predict cells with arrested cell cycle (*e.g.* senescent/quiescent cells), in line with the upregulation of *sin-lncRNA* upon OIS.

While tumor cells are highly proliferative, it has been shown that chemotherapy can induce cellular senescence, which can be linked to the resistance to treatment ^7^. To investigate the potential significance of *sin-lncRNA* in response to cancer therapy, we analyzed the expression of the lncRNA in cisplatin-treated ovarian cancer cells, given their ability to undergo senescence upon treatment with platinum-based agents ^56^. 10 days after cisplatin treatment, COV362 cells displayed a fully senescent phenotype, as confirmed by the expression of senescence markers, such as β-galactosidase and the induction of cell cycle inhibitors and SASP components (Figure 6A-B). Interestingly, we observed a strong increase of *sin-lncRNA* expression in the senescent cells, whereas it was undetectable in untreated cells (Figure 6C). Cisplatin treatment also upregulated C/EBPβ expression (Figure 6D), suggesting that, also in the cancer cells, this transcription factor is responsible for the transcriptional activation of *sin-lncRNA*. As previously shown for senescent fibroblasts, knockdown of *sin-lncRNA* in cisplatin-treated COV362 cells reduced the expression of mRNAs involved in OXPHOS pathway (Figure 6E), which was reflected in alterations in the metabolic functions, as a decreased maximal respiration and increased ECAR (Figure 6F, G). Moreover, knockdown of either *sin-lncRNA* or DLST further decreased cell growth compared to control senescent cells (Figure 6H) indicating that depletion of the senescence-specific lncRNA sensitizes ovarian cancer senescent cells to cisplatin treatment.

**Figure 6.**
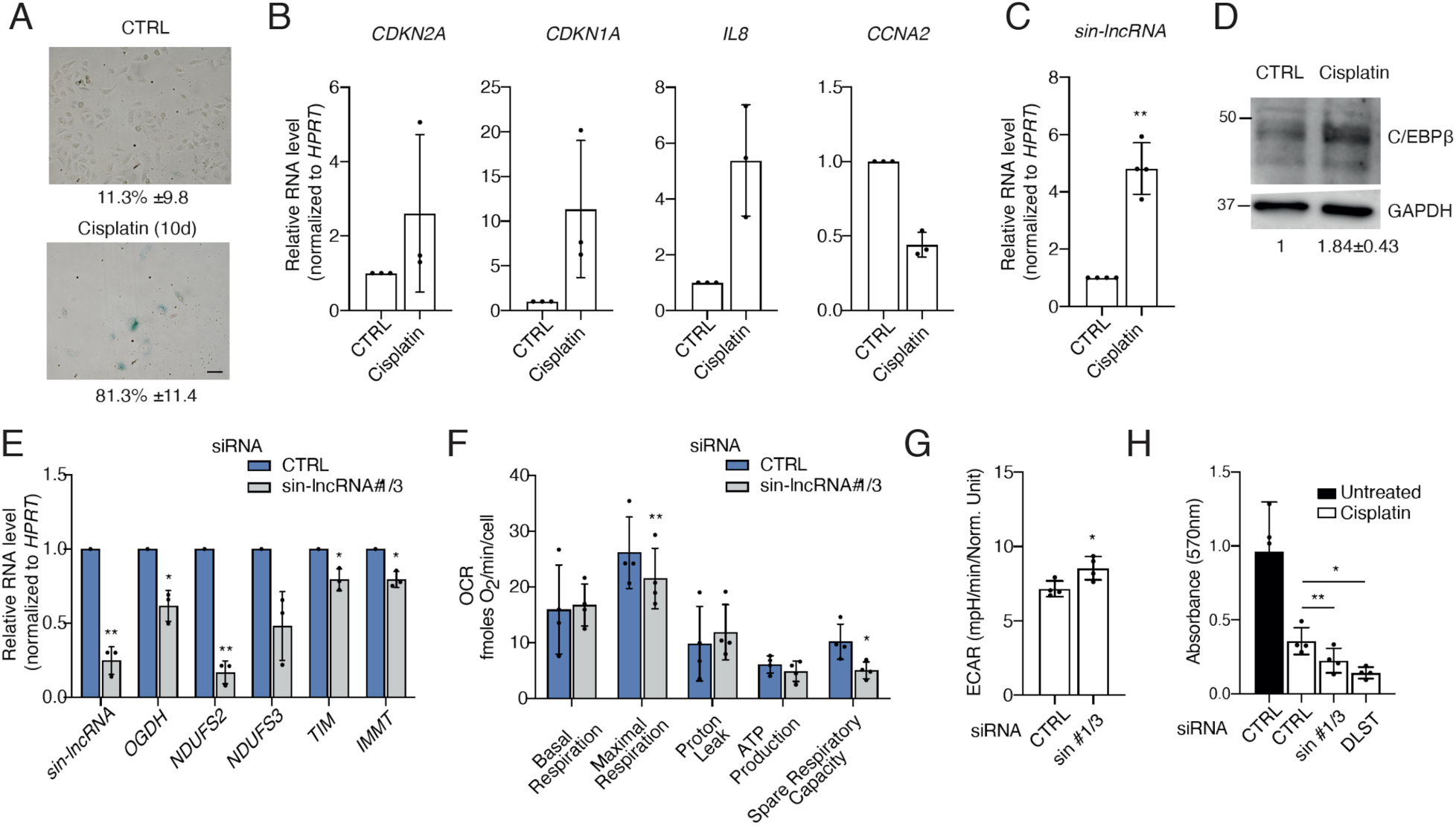
*sin-lncRNA* regulates OXPHOS in ovarian cancer cells. **A)** Representative β-galactosidase images of untreated COV362 cells (upper panel) or treated for 10 days with μM cisplatin. Scale bar: 50 μm. Numbers represent the mean ± s.d. (n=3). **B)** RT-qPCR analysis of senescence markers genes in cisplatin treated COV362 for 10 days compared to untreated controls (set as 1). **C)** RT-qPCR analysis of *sin-lncRNA* in the cells described in B. The graph represents the mean ± s.d. (n=4). One sample *t*-test, \**\*p<*0.01. **D)** Western blot analysis of C/EBPβ protein levels in the cells described in B. Numbers represent the mean ± s.d. (n=3). **E)** RT-qPCR analysis of OXPHOS GSEA pathway genes in control and *sin-lncRNA* depleted (siRNA#1/3) COV362 senescent cells treated with cisplatin for 10 days. **F)** Oxygen Consumption Rate (OCR) of control or *sin-lncRNA* depleted (siRNA#1/3) senescent COV362 cells obtained using the Seahorse XFe96 Analyzer following injections of oligomycin, FCCP, and rotenone/antimycin A, as indicated in the Cell Mito Stress Test. Data is presented as mean ± s.d. (n = 4). Two tailed student’s *t*-test, *\*p<*0.05; \**\*p<*0.01. **G)** Basal extracellular acidification rate (ECAR) measured using the Seahorse XFe96 Analyzer, in the same conditions as in F, and following injections of glucose, oligomycin, and 2-deoxy-D-glucose, as indicated in the Glycolysis Stress Test. Data is presented as mean ± s.d. (n = 4). Two tailed student’s *t*-test, *\*p<*0.05. **H)** Absorbance levels of crystal violet solution measured at 560nm in proliferating or senescent COV362 cells transfected with control, *sin-lncRNA* or DLST siRNA treated for 10 days with 10 μM cisplatin.

Altogether, these data provide evidence that a lncRNA uniquely expressed in senescent cells contributes to the configuration of their metabolic state, also when senescence is induced in cancer cells, thus representing a valuable marker and potential target in aging and cancer.

## DISCUSSION

Although senescence has been traditionally considered a static terminal condition, it is now recognized as a dynamic stepwise process, which involves a plethora of factors that coordinately ensure its correct progress. We have identified a lncRNA that is specifically expressed at later stages of senescence induction to maintain the metabolic balance of these cells. Such precise induction is linked to a timely transcriptional activation mediated by the SASP master regulator C/EBPβ; a transcription factor known to promote the later inflammatory phase of the senescent program ^31^. Remarkably, depletion of *sin-lncRNA* is linked to the activation of some inflammatory signals as well as extracellular matrix remodeling pathways, gene networks that are induced during the second phase of senescence induction. *sin-lncRNA* depletion reinforces the senescence response, altering many of the canonical senescence markers. This is likely a consequence of the augmented ROS production, which feeds forward the senescent phenotype.

Mitochondria are the main source of ROS in the cell ^57^. Interestingly, one of the most deregulated pathways upon *sin-lncRNA* depletion is OXPHOS, which reflects mitochondrial dysfunction and results in high levels of ROS production ^58^. High levels of ROS are critical for onset and maintenance of cellular senescence, although it remains difficult to decipher whether they are the cause or consequence of the senescence process ^59^. Nevertheless, several factors are involved in feedback mechanisms to prevent an uncontrolled ROS increase ^60^. We have found that *sin-lncRNA* interacts with DLST, an enzyme of the TCA cycle, a process that is tightly interconnected to OXPHOS and a main source of ROS ^61^. Interestingly, it has been previously shown that the KGDH complex is sensitive to ROS and involved in ROS production ^62^, and inhibition of this enzyme might be critical in the metabolic deficiency induced by oxidative stress ^63^. DLST depletion in senescent cells mimics *sin-lncRNA* knockdown by exacerbating growth arrest and β-galactosidase staining. Moreover, both *sin-lncRNA* and DLST downregulation result in increased extracellular acidification rate and lactate production, suggesting a shift in the energy production pathway towards glycolysis. This may reflect a mechanism for escaping the restrictions caused by the increased oxidative stress. Interestingly, we have observed different effects when measuring the specific glutamine-derived TCA metabolites by LC-MS. Whereas inhibition of DLST strongly affects different steps of the TCA cycle, sin-lncRNA depletion does not result in such changes. The fact that different enzymes involved in the GABA shunt pathway are found to be overexpressed in these cells, would explain these differences as it could contribute to maintaining TCA cycle activity in *sin-lncRNA* depleted cells. This compensatory mechanism might also explain the fact that the basal respiration rate is not affected in *sin-lncRNA* knockdown cells, while the maximal respiration and the spare capacity are strongly decreased. Previous studies have described similar changes in mitochondrial capacity following silencing of the ACC ADP/ATP carriers, suggesting an important role for AACs in maintaining metabolic spare capacity ^64^. Besides, inhibition of ATP synthase activity has been shown to promote metabolic rewiring to an enhanced aerobic glycolysis and the subsequent production of mitochondrial reactive oxygen species ^37^. Additional transcriptional alterations observed in metabolic genes such as ADP/ATP carriers might contribute to the phenotypic alterations observed upon *sin-lncRNA* knockdown.

Of note, we have identified as a potential *sin-lncRNA* interactor DDX28, which plays a role in OXPHOS and is also known to shuttle from the nucleus to the cytoplasm and mitochondria ^47^. Therefore, we cannot exclude that *sin-lncRNA* might regulate other aspects of metabolism at the interface between cytoplasm and mitochondria or even other subcellular compartments. Interestingly, TCA enzymes have been found to translocate into the nucleus where they have roles in epigenetic regulation ^65–68^. In particular, the KGDH complex which generates succinyl-CoA, has been demonstrated to contribute to histone succinylation affecting expression of cell cycle genes ^52^ and altering metabolic states ^69^. Overexpression of wild type DLST is able to reproduce the effect of *sin-lncRNA* depletion on the transcription of metabolic genes, whereas a mutant which is not able to translocate to the nucleus does not (Figure 5E). Although the mechanism underlying this response needs to be further explored, these results indicate that the two forms have a different impact in transcriptional regulation. Interestingly, we observed a significant overlap between the genes deregulated by the knockdown of *sin-lncRNA* and the H3K79Succ peaks, with key metabolic genes affected ^52^ (Suppl. Figure 6D, E and Fig. 5F), suggesting that *sin-lncRNA* knockdown leads to changes in this histone mark regulating metabolic transcription programs. Thus, while sin-lncRNA expression in senescent cells is relatively high (∼80 molecules per cell), it does not appear sufficient to serve as a structural scaffold for the KGDH complex. Instead, sin-lncRNA likely functions as a signal amplifier, linking mitochondrial processes to nuclear transcriptional programs. The molecular mechanism of interaction between DLST (KGDH complex) and sin-lncRNA, and how it influences the localization of the protein remains to be understood, our observations are in line with the growing evidence that many metabolic enzymes interact with regulatory RNAs ^27, 70^. For instance, recent pre-print work from the Hentze laboratory has shown that RNA is required for proper mitochondrial import of ATP5A1^71^. All this emerging evidence suggests the presence of an undiscovered RNA-protein network with significant implications for regulating cellular energy metabolism.

Platinum-based therapies, currently used as the first line of treatment for patients with ovarian cancer, are known to induce senescence in cancer cells ^72^. Although therapy-induced senescence (TIS) has been considered a desirable outcome of cancer therapy, the activation of the senescence program has been shown to induce the rewiring of cellular metabolism promoting the pro-tumorigenic and pro-metastatic potential of ovarian cancer cells ^73^. Here, we show that *sin-lncRNA* is specifically induced in ovarian cancer cells that undergo senescence and contributes to sustain the metabolic demands to maintain senescence in cancer cells upon drug treatment. Emerging studies link metabolic reprogramming to cancer drug resistance ^74^. Indeed, cisplatin-resistant cells exhibit increased fatty acids (FA) uptake, accompanied by decreased glucose uptake and lipogenesis, indicating reprogramming from glucose to FA dependent anabolic and energy metabolism ^75^. Importantly, other metabolic processes affected by *sin-lncRNA* depletion in senescent fibroblasts seen at the transcriptional level include fatty-acid metabolism. Depletion of *sin-lncRNA* might sensitize ovarian tumor cells to cisplatin by metabolic reprogramming at different levels.

Metabolic enzymes required for the tricarboxylic acid cycle have been shown to affect senescence by regulating p53 ^76^. Importantly, advances in the field of RNA interactome capture have demonstrated that the interplay between metabolic enzymes and RNA is broader than previously considered ^27, 70^. These studies have revealed the RNA binding capacity of many enzymes, although the precise mechanism or the physiological relevance requires deeper characterization. Emerging evidence indicates that lncRNAs may contribute to cellular processes in order to adapt and coordinate mitochondrial function to adjust their activities to the environmental conditions ^27, 77, 78^. We propose that distinct cellular states or stresses that result in metabolic changes involve the function of ncRNAs that contribute to the reconfiguration of the cell metabolism. The exquisite specificity of *sin-lncRNA* in senescence highlights its potential role as a therapeutic target. This discovery paves the way for better comprehension of therapy-induced senescence and its associated implications.

## MATERIALS AND METHODS

### Cell culture, retroviral infection and treatments

IMR90 lung fibroblasts and HEK-293T cells were purchased from ATCC. TIG3 hTERT pMSCV-ER:ΔBRAF lung fibroblasts ^33^ were kindly provided by Dr. Lund’s laboratory (BRIC – Copenhagen, Denmark). All cell lines were cultured in DMEM medium (GIBCO), supplemented with 10% fetal bovine serum (GIBCO) and 1x penicillin/streptomycin (Lonza). Cells were maintained at 37°C in the presence of 5% CO2 and tested for mycoplasma contamination regularly, using the MycoAlert Mycoplasma Detection Kit (Lonza).

To generate OIS cell systems, viruses were first produced in HEK-293T cells transfected with 12μg of either pLNC-ER:RAS or pLNCX (Empty) vectors, 6μg of gag-pol plasmid, 3μg of pVSVG vector and 1μg of pMAX-GFP plasmid (to check transfection efficiency). Transfection reaction was carried out in opti-MEM medium (GIBCO) using Lipofectamine 2000 (Invitrogen), following manufacturer’s instructions. 48 hours after transfection, filtered supernatant supplemented with 4μg/ml polybrene (Santa Cruz), was used to transduce low passage IMR90. Cells were selected with Neomycin-G418 (Sigma) at a final concentration of 400μg/ml for at least one week.

For senescence induction studies, IMR90 ER:RAS or TIG3-hTERT ΔER:BRAF fibroblasts have been treated with 200nM of 4-hydroxy-tamoxifen (4OHT) for three days unless specified otherwise in the text. When longer time courses were needed, medium was replaced on day 3. Only normal fibroblasts cell lines with less than 20-25 passages were used for experiments.

COV362 cells were obtained from SIGMA, kindly provided by Dr. Beatriz Tavira (CIMA) and cultured in DMEM medium (GIBCO), supplemented with 10% fetal bovine serum (GIBCO) and 1x penicillin/streptomycin (Lonza). For senescence induction the cells were treated with 10 μM cisplatin for 10 days.

N-Acetyl-L-cysteine (NAC, SIGMA) was added at 5 mM 1h prior 4-OHT treatment and for the duration of the experiment.

### DLST WT and DLST NLS mutant –overexpressing cell lines

To generate lentiviral constructs, we cloned the full-length *DLST* sequence amplified from cDNA into the inducible vector pLVX-TetOne Puro Vector (Clontech). For the mutant form, we used as a template a plasmid sintesized by Genescript. We used In-Fusion HD Cloning designing primers according to the manufacturer’s indications (see primer sequences in Suppl. Table 5).

For the lentiviral production, HEK293T cells were transfected using Lipofectamine 2000 (Life Technologies) with 7μg of pLTVX-WT-DLST or pLTVX-Mut-DLST (R224A/K226E), together with 6μg of VsVg and 5μg Pax8 viral plasmids. After 48h, supernatants containing the viral particles were filtered through a 0.45-μm filter. Confluent IMR90 ER:RAS cells were then infected with one-third of the viral supernatant and 8μg/μl polybrene. Twenty-four hours post infection, the cells were selected with 1μg/μl puromycin. Cells were maintained in tetracycline-free tested serum (Clontech). For the expression of the protein 100ng/ml doxycicline was used for 72h.

### Irradiation

To induce senescence by ionizing radiation, IMR90 cells were γ-irradiated at 5Gy and harvested at the indicated times. Radiation was delivered at 180 cGy/min using a Siemens Oncor Impression Plus linear accelerator equipped with 6MeV X rays.

### RNAi studies

For siRNA studies, cells were transfected once with a final concentration of 40nM siRNA using Lipofectamine 2000 (Invitrogen) in opti-MEM medium (GIBCO), unless differently stated in the text. Scrambled siRNA (siRNA CTRL in the text) was used as transfection control. All siRNAs employed in this study were designed using BLOCK-iTTM RNAi Designer (https://rnaidesigner.thermofisher.com/rnaiexpress/) and purchased from Sigma, except C/EBPβ siRNA, a kind gift of Dr. Aragón (CIMA – Pamplona, Spain).

To induce senescence in transfected cells, the transfection medium was replaced after 6 hours with complete medium containing 200nM 4OHT. When a senescence induction longer than 3 days was needed, transfected cells were trypsinized and re-plated to perform multiple analyses in parallel. All siRNAs used in this study are listed in Suppl. Table 5.

### CRISPR-Cas9 gene editing and CRISPR clonal cell lines generation

The CRISPR Design Tool from the Zhang Lab (http://crispr.mit.edu/) was used to find suitable target sites for the Streptococcus pyogenes Cas9 (SpCas9) in *sin-lncRNA* genomic region. Specific guide RNAs (sgRNA_up and sgRNA_dw, see Suppl. Table 5) were cloned into the lentiviral plasmid lentiCRISPRv2 carrying spCas9 gene and puromycin resistance, a kind gift from Dr. Agami’s lab (NKI – Amsterdam, Netherlands) and originally generated in Dr. Zhang’s lab (for details, see plasmid reference #5296113 on Addgene website). LentiCRISPR vectors carrying the two independent sgRNAs were separately transfected in HEK-293T cells to generate viral particles, as described above. LentiCRISPR viruses were mixed (ratio 1:1) and used to transduce TIG3-hTERT ΔER:BRAF fibroblasts, then selected with 1μg/ml of puromycin. Single puromycin-resistant TIG3 cells were sorted in 96-well plates, using untreated TIG3 fibroblasts as feeder cells, which were then eliminated by a second round of puromycin selection (0.5μg/ml). Genomic DNA was isolated from puromycin-resistant clones using QuickExtract DNA extraction solution (Lucigen), PCR amplified and tested on agarose gel for accurate deletion.

### Cell proliferation assays

For crystal violet assay upon OIS, 1-2×10^4^ untreated or transfected cells were plated in 6-well plates and senescence induction was prolonged for 5 days in 4-OHT medium. Cells were fixed with 0.5% glutaraldehyde for 15 min and stained with 0.1% Crystal Violet solution (Sigma) for 30 min. To quantify cell density, cells were incubated with 500μl of 10% acetic acid (Sigma) and collected in ELISA plates. Absorbance was measured by spectrophotometry at 570 nm in a SPECTROstar Nano equipment.

For BrdU incorporation assay, cells were labeled with 50 μM BrdU solution (BD Pharmingen) for 16-18 hours. In case of senescent cells, OIS was previously induced for 5 days before starting BrdU assay. Then, cells were harvested by trypsinization and BrdU staining was performed using BrdU Flow Kits (BD Pharmingen), according to manufacturer’s instructions. Amount of BrdU incorporation was measured by flow cytometry using a FACSCalibur Cell Analyzer (BD Bioscience).

### Apoptosis staining

Apoptosis was measured using BD Pharmingen™ PE Annexin V Apoptosis Detection Kit I (BD Bioscience). Briefly, cells were washed twice with cold PBS and then resuspend cells in 1X Binding Buffer at a concentration of 1 x 10^6 cells/ml. 1 x 10^5 cells were stained with 5 μl of PE Annexin V and 5 μl 7-AAD. After 15 mins incubation at RT in the dark. 400 μl of 1X Binding Buffer was added to each tube. Apoptosis levels were measured by flow cytometry using a FACSCalibur Cell Analyzer (BD Bioscience).

### Senescence Associated galactosidase (SA-β-gal) staining

To detect SA-β-gal activity, 1×10^3^ of untreated or transfected cells were plated in 6-well plates and senescence was induced for 8 days upon 4-OHT treatment. Cells were fixed with 0.5% glutaraldehyde solution for 15 min at RT and washed with PBS containing 1mM MgCl2 (pH 6.0). Then, cells were incubated overnight at 37°C in X-gal staining solution (1mg/ml X-gal [Thermo Fisher], 0.12 mM K3Fe(CN)6 [Merk], 0.12 mM K4Fe(CN)6 [Merk], PBS/MgCl2 [pH 6.0]). Cells were washed and percentage of blue cells was counted manually using an inverted microscope.

### ROS/RNS measurement assay

Senescent IMR90 ER:RAS fibroblasts transfected with *sin-lncRNA* targeting or control siRNAs were collected by trypsinization and re-suspended at 0.5-1×10^6^ cells/ml in Hypotonic Buffer (10mM Tris HCl [pH 7.5], 10mM NaCl, 1.5mM MgCl2). To facilitate membrane lysis, cells were incubated for 10 min at 4°C and sonicated for 5 cycles (30’’ON-30’’OFF) in a Bioruptor sonication device. Cell extract was cleared by centrifugation and assayed for ROS/RNS accumulation by fluorescence-based approach using OxiSelect in vitro ROS/RNS Assay Kit (Cell Biolabs), following manufacturer’s instructions. The amount of fluorescent oxidized probe (2’, 7’ –dichlorodihydrofluorescein or DCF) was measured with a fluorescence plate reader at 480nm excitation / 530nM emission. Data were interpolated to a DCF standard curve to obtain absolute values.

### Measurement of oxygen consumption and extracellular acidification

Cellular OCR was measured using a Seahorse XF Cell Mito Stress Test and a Seahorse Bioanalyzer XFe96 (Agilent Technologies) according to the manufacturer’s standard protocol. Briefly, 200,000 IMR90 cells were reverse transfected with the corresponding siRNAs in 6 well plates. After 24h cells were treated with 500 nM 4-OHT. After 3 days of treatment, cells were trypsinized and seeded on Seahorse XF96 Cell Culture microplates to confluence (10000 cells/well) and were subjected to metabolic profiling the next day. For Mito stress tests, cells were cultured for 1 h in a CO_2_-free incubator at 37 °C. Oxygen consumption rates (OCR) were monitored at basal conditions and, after sequential injections of 1μM oligomycin to block the mitochondrial ATP synthase, 1.5μM FCCP to uncouple oxidative phosphorylation, 1μM antimycin A and 1μM rotenone were used to fully inhibit mitochondrial respiration and 50 mM 2-deoxyglucose (2-dG) to block glycolysis. Two separate measurements of OCR and extracellular acidification rate (ECAR) from 4 independent experiments were taken after the addition of each inhibitor. Samples were normalized based on the protein concentration. All reagents were purchased from Agilent Technologies.

### Lactate measurements

IMR90 ER:RAS fibroblasts were transfected with *sin-lncRNA* targeting or control siRNAs and treated with 4-OHT for 6 days. The supernatant was harvested and centrifuged for 5 min at 500 g. Lactate levels were measured using Cobas C311 Analyzer with LACT2 kits (Roche Diagnostics GmbH). Human Precinorm y Precipath were used as controls.

### Liquid Chromatographic –Mass Spectrometry (LC-MS) metabolomics

#### Isotope labelling

6-days senescent IMR90 ER:RAS fibroblasts transfected with siRNA control or siRNA against *sin-lncRNA* or DLST were incubated with glutamine-free DMEM supplemented with 4mM L-^13^C_5_-Glutamine (Cambridge Isotopes Laboratories). After 6 hours the cells were washed with PBS 1x and snap-freeze in liquid nitrogen.

#### Sample preparation and LC/MS methods

LCMS was performed at the Metabolomics Platform at CIC bioGUNE (Donostia, Spain). Cells from three wells (1E6 cells/well) were pooled to obtain one LCMS sample. Therefore, cells were extracted with 200 μL icecold extraction liquid per well. The volume from one well was transferred to the next and finally the resulting 600 μL volume was passed over the three wells two times. The extraction liquid consisted of a mixture of ice-cold methanol/water (50/50 %v/v). Subsequently 400 μL of the cell homogenate plus 400μL of chloroform was transferred to a new aliquot and shaken at 1400 rpm for 60 minutes at 4 °C. Next the aliquots were centrifuged for 30 minutes at 14000 rpm at 4 °C. 250μL of the aqueous phase was transferred to a fresh aliquot. The chilled supernatants were evaporated with a speedvac in approximately 2h. The resulting pellets were resuspended in 150 μL water/MeOH (75/25 %v/v). Samples were measured with a UPLC system (Acquity, Waters Inc., Manchester, UK) coupled to a Time-of-Flight mass spectrometer (ToF MS, SYNAPT G2S, Waters Inc.). A 2.1 x 100 mm, 1.7 μm Phenyl-Hexyl column (Waters Inc.), thermostated at 40 °C, was used to separate the analytes before entering the MS. Mobile phase solvent A (aqueous phase) consisted of 99.5% water and 0.5% FA while solvent B (organic phase) consisted of 99.5% MeCN and 0.5% FA. In order to obtain a good separation of the analytes the following gradient was used: from 98% A to 0% A in 2 minutes in curved gradient (#8, as defined by Waters), constant at 0% A for 1 minute, back to 98% A in 0.2 minutes. The flow rate was 0.250 mL/min and the injection volume was 3 μL. After every 6 injections, a sample was injected. The MS was operated in negative electrospray ionization in full scan mode. The cone voltage was 10 V, cone offset was 50V and capillary voltage was 400 V. Source temperature was set to 120 °C and capillary temperature to 500 °C. The flow of the cone, desolvation and nebulizer gas (nitrogen) were set to 5 L/h, 6 L/h and 850 L/h, respectively. A 2 ng/mL leucine-enkephalin solution in water/acetonitrile/formic acid (49.9/50/0.1 %v/v/v) was infused at 10 μL/min and used for a lock mass which was measured each 36 seconds for 0.5 seconds. Spectral peaks were automatically corrected for deviations in the lock mass. Extracted ion traces for relevant analytes were obtained in a 10 mDa window in their expected *m/z*-channels. These traces were subsequently smoothed and peak areas integrated with TargetLynx software. Signals of labeled analytes were corrected for naturally occurring isotopes. Since we also had unlabeled experiments, for some metabolites we could determine the naturally occurring isotope-distribution empirically. This was necessary for the first and second isotopomers since correcting with theoretical values did not eliminate these isotopes from unlabeled experiments. This is probably because signals are non-linear with respect to ion-counts, *i.e.* higher ion-counts tend to have *relatively* lower signals than lower ion-counts due to saturation of the detector. The calculated raw signals were adjusted by median fold-change (MFC) adjustment. This is a robust adjustment factor for global variations in signal due to e.g. difference in tissue amounts, signal drift or evaporation. The MFC is based on the total amount of detected mass spectrometric features (unique retention time/mass pairs). The calculations and performance of the MFC adjustment factors are described in the following publications ^79, 80^.

### Immunofluorescence and RNA fluorescence in situ hybridization (RNA-FISH)

For fluorescence-based studies, IMR90 ER:RAS cells were collected by trypsinization after required treatments (as specified in the text) and re-plated on top of cover glasses placed on clean 6-well plates. The following day, cells were washed and fixed for 15 min at room temperature (RT) using 3% methanol-free formaldehyde solution (Thermo Fisher). For immunofluorescence studies, fixed cells were washed (PBS/0.5% NP-40) and incubated first with block solution (10% FBS in wash buffer) for 20 min and then with primary antibody diluted in block solution for 1 hour at RT. Antibody solution was washed out and cells were incubated with fluorophore-conjugated secondary antibody for 30 min. After extensive washes, cover glasses were mounted on microscope slides using DAPI-containing mounting solution (Palex Medicals) and pictures were collected with an automated optical microscope running Zen 2 core imaging software (Zeiss), according to standard procedures. For RNA-ISH studies, fluorescein-labeled Locked Nucleic Acid (LNA) DNA probes were designed and synthesized by Exiqon and were hybridized according to manufacturer’s protocol with some modifications. Briefly, fixed cells were first incubated with 70% ethanol for 1 hour and then with Acetylation Buffer (0.1M Triethanol Amine, 0.5% (v/v) Acetic Anhydride) for 30 min at RT. To avoid non-specific probe binding, cells were incubated with Hybridization buffer (10% dextran sulfate, 50% formamide, 2x saline-sodium citrate [SSC] buffer) for 1 hour at 55°C. Specific LNA probes were denatured at 92°C for 4 min and then mixed with Hybridization buffer at a final concentration of 20nM. Hybridization was carried out overnight at 55°C. The following day, probes residues were eliminated using 2x SSC wash buffer and fixed cells were treated with 3% Hydrogen Peroxide for 30 min at RT. For fluorescein (FAM) detection, cells were first incubated 1 hour with Blocking Buffer (10% heat-inactivated Goat Serum, 0.5% Blocking Reagent [Roche] in PBS-0.5% Tween) and then 1 hour with 1.5U/ml αFAM-POD antibody (Roche) diluted in Blocking Buffer. After extensive washes, fluorescent signal was developed by 10 min incubation with TSA-Cy3 solution (Perkin Elmer). Residues were eliminated through stringent 4x SSC washing and slides were prepared for microscope detection using DAPI mounting solution, as mentioned above.

Slides were imaged on a fluorescence inverted microscope using a 63x objective. At least 10 optical stacks were acquired per image for DAPI (nuclear stain) and for the fluorescence channel corresponding to the RNA FISH probe (Cy3). Image processing was performed using Zen software (version 2.3 – Zeiss) and for each condition, signal was adjusted on a sample incubated with no probes and used as control for background detection. ImageJ software was employed for stacks deconvolution.

### Protein extraction and immunoblot analysis

Cells were lysed for 15 min in rotation at 4°C using RIPA buffer (150mM NaCl, 25mM Tris HCl [pH 7.5], 2mM EDTA, 0.1% sodium deoxycholate [Na-DOC], 0.1% SDS, 1% Triton X-100) supplemented with 1x cOmplete Protease Inhibitor Cocktail [Roche]. For detection of phosphoproteins, 1x PhosSTOP phosphatase inhibitor [Roche] was freshly added to RIPA solution. Lysed cells were centrifuged at max speed for 10 min at 4°C and the insoluble pellet was discarded. Protein concentration was estimated by Pierce BCA Protein Assay Kit, using BSA curve as reference and according to manufacturer’s instructions. Proteins were separated on denaturing SDS-PAGE gels and transferred to a nitrocellulose membrane [Biorad] following standard procedures. Membranes were blocked using skim milk or BSA (VWR) and probed first for primary and then for HRP-conjugated secondary antibody. Western Lightening ECL-Plus (Perkin Elmer) was employed for chemiluminescence detection of proteins. The list of antibodies used in this study can be found in Suppl. Table 5.

### Cellular Fractionation

IMR90 cells were reverse transfected with control or siRNA 1/3 *sin-lncRNA*. The day after transfection cells were treated with 500 nM 4-OHT for 5 days. Cells were scrapped in cold PBS and centrifuge 5 min 0.2g. Pellets were resuspended in NIB buffer (1.28 M Sucrose, 40 mM Tris pH 7.4, 20 mM MgCl2, 4% Triton X-100), incubated on ice for 20 min and centrifuged 15 min at 2500rpm. The supernatant was kept as the cytosolic fraction and further cleaned by an extra centrifugation step. Nuclear pellets were washed twice with NWB (40 mM Tris pH 7.4, 20 mM MgCl_2_) by centrifugation 15 min at 2500 rpm. RIPA (150 mM NaCL, 25 mM Tris pH 8, 0.1% SDS, 0.1 % Na-Deoxycholate, 2 mM EDTA, 1% Triton X-100) buffer was used to lyse the nuclei. After 10 min centrifugation at max speed the supernatant was isolated as the nuclear fraction. Immunoblot analysis was performed as described above.

### Mitochondria isolation

10 cm plates of senescent cells were incubated with 200nM of MitoTracker for 30 min (Invitrogen). Cells were collected by trypsinization and resuspended in 1ml of PBS. Cells were lysed using by 20 strokes up and down using a 2 ml cell douncer. After 5 min centrigfugation at 700g at 4°C the supernatant was collected. Mitochondria were pellet by centrifugation at 12,000g for 5 min at 4°C, washed with PBS once and resuspended in 200 ml of filtered PBS. Mitochondria were purified by FACS sorter using Flow Cytometry Size Calibration Kit (Invitrogen).

### Native RNA immunoprecipitation

20×10^6^ cells were collected in cold PBS by scraping and lysed in RIP buffer (25mM Tris-HCl pH 7.4, 150mM KCl, 5mM EDTA, 0.5% Nonidet P-40, 20Uml^−1^ RNAsin (Promega), 0,5mM dithiothreitol (DTT) and 1× protease inhibitor mixture Complete (Roche)). After centrifugation, the samples were precleared with Dynabeads Protein A (Invitrogen) for 1h. One per cent of the sample was used as the input control and the remaining extracts were incubated with 10 μg DLST antibody (Bethyl) or IgG at 4°C overnight. RNA–antibody complexes were collected by incubation with Dynabeads Protein A. After extensive washing in RIP buffer (25mM Tris-HCl pH 7.4, 150mM KCl, 5mM EDTA, 0.5% Nonidet P-40, 20Uml^−1^ RNAsin (Promega), 0,5mM dithiothreitol (DTT) and 1× protease inhibitor mixture Complete (Roche)), 1/5 of the sample was used for western blotting control and the remaining for RNA extraction and qRT–PCR analysis, as described below.

### *In vitro* RNA pulldown

RNA pulldown was performed according to ^81^. Briefly, biotin-labeled *sin-lncRNA* and control RNA (cloned in a pcDNA3.1(+) plasmid) were generated in vitro using T7 or T3 RNA polymerase, respectively (Applied Biosystems). The fragments used for the mapping were in vitro transcribed from PCR products carrying the sequence of the T7 promoter in the forward primer (see Suppl. Table 5), Biotinylated RNAs were treated with RNase-free DNase I (Promega), purified on G-50 MicroSpin columns (GE Healthcare) and heated to 65°C for 10 min and cooled down slowly to 4°C to allow proper RNA folding. 10×10^6^ IMR90 ER:RAS were collected after 6 days of senescence induction and total protein extracts were prepared using RIPA buffer. Cell extracts were then incubated with 10 μg of biotinylated RNA in Buffer A (150mM KCl, 25mM Tris pH 7.4, 5mM EDTA, 0.5mM DTT, 0.5% NP40) supplemented with fresh protease inhibitors and yeast tRNA (0.1μg/ml) and rotated overnight at 4C. Dynabeads MyOne Streptavidin T1 beads (Invitrogen) were used to isolate RNA-protein complexes. Beads were washed 5 times in buffer A (150 mM KCl, 25 mM Tris pH 7.4, 5 mM EDTA, 0.5 mM DTT, 0.5 % NP40, 1x protease inhibitor cocktail (Roche), 1 mM PMSF, 100 U/ml SUPERASin) and boiled in 2X Laemmli loading buffer, and retrieved proteins were loaded in a NuPAGE 4%–12% bis-Tris gel (Invitrogen). Gel was stained with the SilverQuest Silver Staining Kit (Thermo Fisher) to identify differential bands comparing *sin-lncRNA* and control RNA samples. We isolated a band of ∼50-55 kDa unique for *sin-lncRNA* (Suppl. Figure 5). Unique bands were cut and submitted for Mass Spectrometry (MS) analysis to the Taplin MS Facility (Harvard University).

For the mapping of the fragments, the pull down was done as indicated above but cisplatin-induced senescence COV362 cells were used.

### Chromatin Immunoprecipitation (ChIP)

10×10^6^ cells were crosslinked with 1% formaldehyde for 10 minutes at RT and nuclei were isolated with cell lysis buffer (5mM Tris pH8.0, 85mM KCl, 0.5% NP40, supplemented with fresh 0,5mM DTT and 1× protease inhibitor mixture Complete (Roche)). Nuclear pellet was then resuspended in RIPA buffer (1X PBS, 1% NP40, 0.5% Na-deoxycholate, 0.1% SDS, supplemented with 1× protease inhibitor mixture Complete (Roche)) and chromatin was sheared to a range of 200-500 bp by using a Bioruptor instrument (Diagenode) with the following settings: 8 cycles LOW, 30’’ON/30’’OFF. Sonicated chromatin was incubated overnight with 6μg of C/EBPβ, H3K79Succ or IgG antibody (listed in Suppl. Table 5) followed by 2h incubation with Dynabeads Protein G beads (Invitrogen). After five washes with LiCl buffer (100mM Tris pH 7.5, 500mM LiCl, 1% NP40, 1% Na-deoxycholate) and TE buffer, DNA was reverse crosslinked by incubating beads in Elution buffer (1% SDS, 0.1M NaHCO3 and Proteinase K (NEB)). C/EBPβ and H3K79Succ binding was analyzed by qPCR using specific primers (see Suppl. Table 5).

### RNA extraction and RT-qPCR analysis

Total RNA was isolated using TRI reagent (Sigma), treated with DNase I (Invitrogen) and reverse-transcribed using the High-Capacity cDNA Reverse Transcription Kit (Applied Biosystem) with random hexamer primers following manufacturer’s instructions. The obtained cDNA was analyzed by quantitative PCR using iTaq Universal SYBR Green supermix (Bio-Rad) in a ViiATM 7 Real-Time PCR System machine (Thermo-Fisher). All reactions were performed in triplicate or quadruplicate and HPRT1 (Hypoxanthine Phosphoribosyl transferase 1) RNA levels were used for normalization, unless specified otherwise in figure legend.

To calculate the number of *sin-lncRNA* molecules per cell, *sin-lncRNA* cloned in pcDNA3.1 vector was in vitro transcribed, quantified and converted to number of copies based on its predicted molecular weight. Dilutions of this transcript were retro-transcribed and used as standard curve for qPCR analysis along with cDNA obtained from a precise number of IMR90 proliferating or senescent cells. All qPCR primer sequences are provided in Suppl. Table 5.

### RNA sequencing and data analysis

Raw data of Poly-A+ RNA-seq from proliferating and senescent IMR90 fibroblasts were kindly provided by Dr. Gil’s laboratory (Imperial College London, UK) before their publication and are now available at the Gene Expression Omnibus under the accession number GSE61130 ^28^. For RNA-seq of IMR90 ER:RAS fibroblasts treated with sin-lncRNA or control siRNA, cells were transfected and 4OHT-treated for 3 days, as described above. Total RNA from samples prepared in biological triplicate was isolated with TRI reagent, purified and treated with DNAse I using RNAeasy mini-Kit (Qiagen). After quality evaluation by Experion kit and quantification by Qubit, 1μg of RNA was used for library preparation and sequenced on Illumina HiSeq 2000 (33×10^6^ reads per sample, 50bp single-end sequencing modality). Sequenced reads from three different replicates were aligned to hg19 genome using the BOWTIE2 algorithm (v.2.1.0) was used to quantify the number of reads in annotated genes ^82, 83^ To identify enriched molecular pathways associated with differences in gene expression, Gene Set Enrichment Analysis (GSEA, Broad Institute, http://www.broadinstitute.org/gsea/index.jsp) was performed using the Hallmark gene sets in the Human MSigDB database ^84^. Enriched pathways were selected with Normalized Enrichment Score (NES) >±1 and adj.Pvalue<1e^-4^.

### *Sin-lncRNA* expression analysis in tumor samples

TCGA RNA-seq expression data was retrieved using R bioconductor library TCGAbiolinks, and represented using R.

### Statistical analysis

Unless specified otherwise, experimental data were represented as mean ± standard deviation of at least three biological replicates (unless otherwise stated in figure legends) and significance was determined by two-tailed unpaired or Student’s t-test or one sample using GraphPad software. Significant P values were summarized as follows: not significant (ns); p-value<0.05 (*); p-value<0.01 (**); p-value<0.001 (***).

## DATA AVAILABILITY

RNA-seq from proliferating and senescent IMR90 fibroblasts were retrieved from public data (GSE61130) ^28^. RNA-seq from replicative senescence mitoplasts were retrieved from public data (GSE73458) ^46^. H3K79Succ peaks were retrieved from public data (GSE97994) ^52^.

RNA-seq of *sin-lncRNA* KD and control senescent cells have been deposited in NCBI’s Gene Expression Omnibus and are accessible through Gene Expression Omnibus (GEO)^85^ repository under the accession number GSE241620 (https://www.ncbi.nlm.nih.gov/geo/query/acc.cgi?acc=GSE241620) with the reviewers token mvylowqmjtotnwx. The mass spectrometry data have been deposited to the ProteomeXchange Consortium via the PRIDE ^86^ partner repository with the dataset identifier PXD044718 (Username, Password: 8egDSySE).

## ACKNOWLEDGEMENTS

We thank Eva Santamaría for her input and help with the seahorse experiments. We thank Beatriz Tavira (CIMA, Universidad de Navarra) for providing the COD362 cells. We thank Sebastiaan Martijn Van Liempd and Juan Manuel Falcón (Metabolomic Platform, CIC bioGUNE) and Lorea Válcarcel (UPV) and Arkaitz Carracerdo (CIC bioGUNE) for their input on the metabolic experiments. We thank Victor Segura and Eli Guruceaga, for the guilt-by-association analysis. We thank Jesus Gil (MRC, London) for his input. We acknowledge the ENCODE project and The Cancer Genome Atlas database for their valuable data sets. We thank the rest of the Huarte lab for contributing with helpful discussions. The work was supported by Project PID2020-113683GB-I00 financed by MICIU/AEI /10.13039/501100011033; La Caixa Foundation [LCF/PR/HR21/00176], Worldwide Cancer Research grant 25-0335 and Ayuda Proyectos Generales AECC 2024. M.M was funded by a MSCA fellowship (HORIZON-MSCA-2021-PF-01: 101066499) and currently by Ministerio de Ciencia e Inovación through Programa Ramón y Cajal (RYC2022-036819-I). and currently by Programa Ramón y Cajal (RYC2022-036819-I). L.P is supported by Instituto de Salud Carlos III (Ministerio de Ciencia e Innovación) through a Miguel Servet contract (CP22/00005), co-funded by the European Union.

## AUTHORS CONTRIBUTION

M.H., M.M. and E.G. conceived the study. E.G, M.M and M.H. designed the experiments, analyzed the data and wrote the manuscript with the comments of co-authors. E.G., F.M., J.G. and M.M. performed the experiments. E. G, J.M.F and A.A. performed bioinformatic analyses. L.P. provided interpretation and made figures of the metabolomics experiment. N.H. contributed to the original set-up of the project.

## FIGURES

**Suppl. Figure 1.**
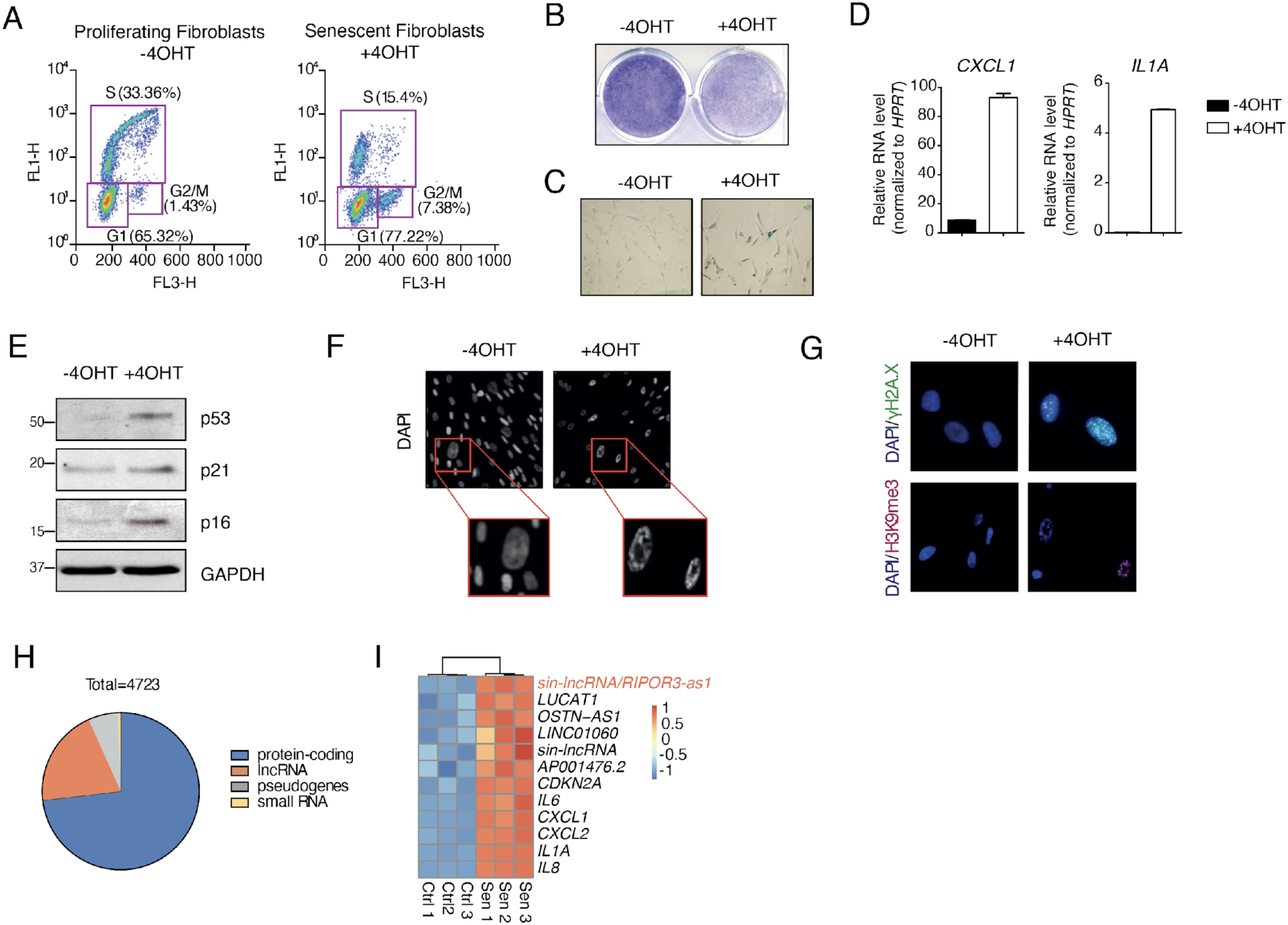
**A**) Change in proliferation rate comparing proliferating and senescent IMR90 ER:RAS fibroblasts, as measured by BrdU assay. **B)** Representative images of crystal violet staining in untreated or 4-OHT treated cells for 5 days. **C)** Representative images of the β-galactosidase staining in untreated cells or treated with 4-OHT for 8 days. **D)** qRT-PCR analysis of CXCL1 and IL1A upregulation in untreated or 4-OHT treated cells for 5 days. **E)** Western blot analysis of senescence markers p53, p21 and p16 in untreated or 4-OHT treated cells for 5 days. **F)** DAPI staining and **G)** Immunostaining images of gH2A.X, H3K96m3 and DAPI of untreated or 4OHT treated IMR90 ER:RAS for 5 days. **H)** Pie chart representation of the number of biotype distribution of transcripts deregulated during OIS (log_2_-fold change >±1, adj.Pvalue <0.05). **I)** Heat map showing relative expression of senescence markers and lncRNAs deregulated in IMR90 ER:RAS untreated or treated with 4-OHT for 8 days.

**Suppl. Figure 2.**
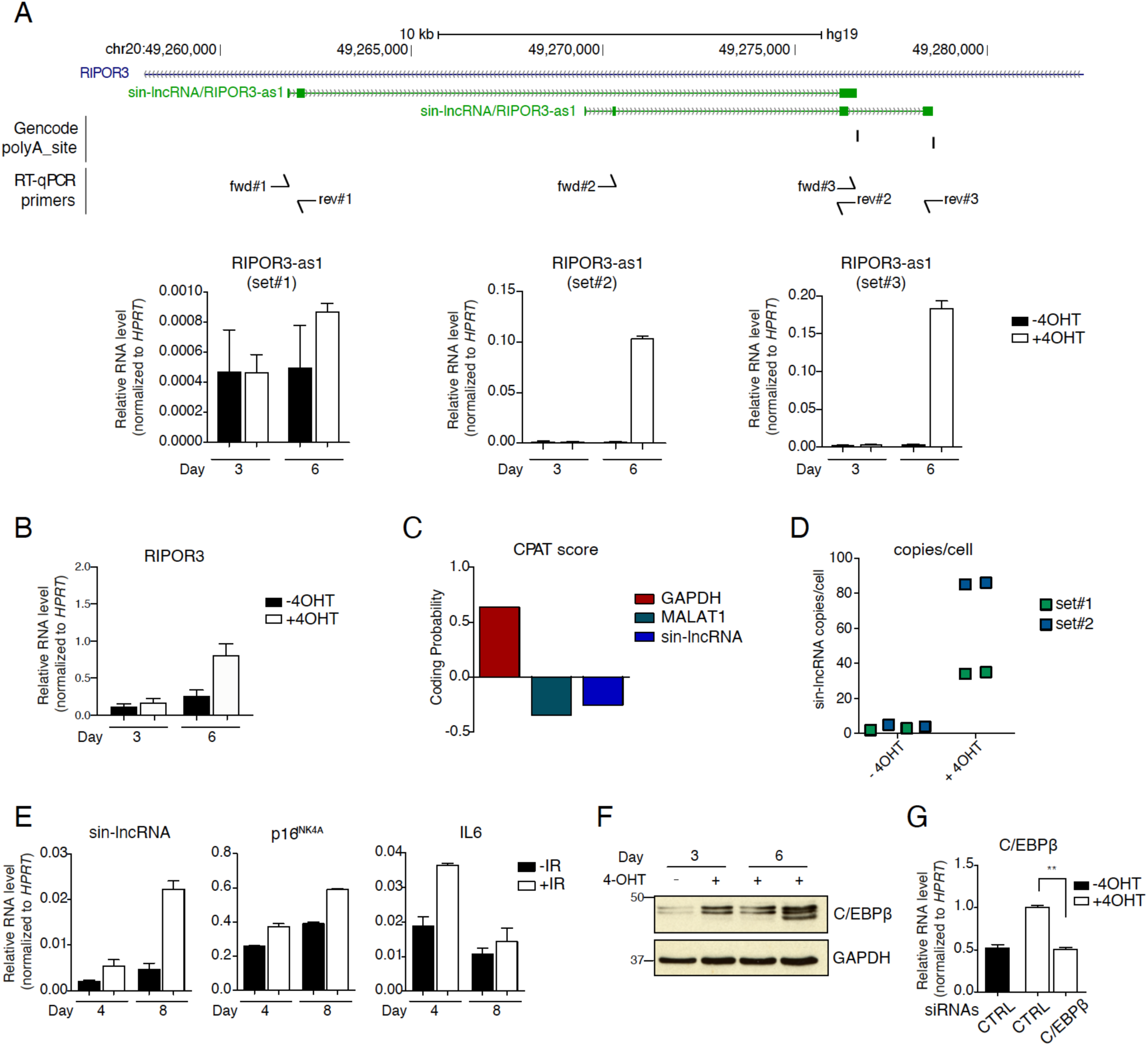
**A**) Genomic snapshot of *sin-lncRNA* (RIPOR3-as) locus PolyA sites annotated by Genecode v19 are indicated as well as the position of qPCR primers used to amplify the different isoforms. Bottom panel, qRT-PCR analysis of the different annotated isoforms of *sin-lncRNA* (RIPOR3-as) using different set of primers (set1-3) in a time course of 3 and 6 days of senescence induction in IMR90 ER:RAS cells. **B)** qRT-PCR analysis of protein-coding gene RIPOR3 in a time course of 3 and 6 days of senescence induction in IMR90 ER:RAS cells. **C)** CPAT software was interrogated to determine the coding probability of *sin-lncRNA*. GAPDH mRNA and MALAT1 lncRNA were used as coding and noncoding references, respectively. **D)** Number of copies per cell of *sin-lncRNA* in control untreated cells and in 4-OHT-treated cells measured by qRT–PCR using dilutions of a pcDNA3.1-*sin-lncRNA* plasmid as standard curve. **E)** RT-qPCR analysis showing *sin-lncRNA* upregulation upon treatment with γ-irradiation at 5Gy in IMR90 fibroblasts. *p16* and *IL6* levels served as positive controls. **F)** Western blot analysis of C/EBPβ of IMR90 ER:RAS cells treated or not with 4OHT for 3 or 6 days. **G)** qRT-PCR analysis of the levels of C/EBPβ in control and C/EBPβ-depleted senescent IMR90 ER:RAS cells

**Suppl. Figure 3.**
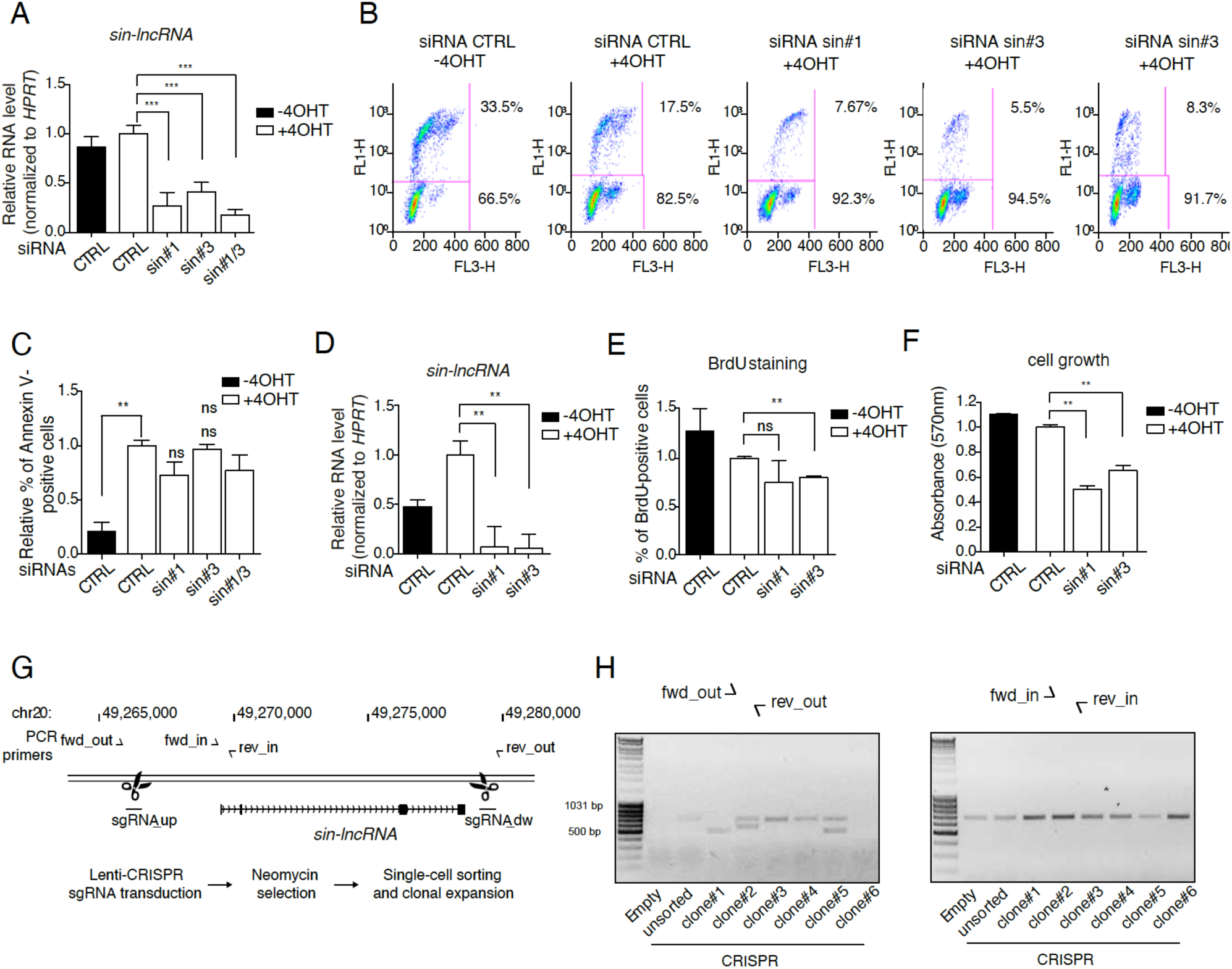
**A**) RT-qPCR analysis of control and *sin-lncRNA* depleted IMR90 ER:RAS cells after 3 days of senescence induction. Proliferating control cells were used as further control. Primer sets #2 (Suppl. Figure 2A) were used to amplify *sin-lncRNA*. Values were normalized to senescent control cells. **B)** BrdU analysis measuring the proliferation of control and *sin-lncRNA* depleted IMR90 ER:RAS cells after 5 days of senescence induction. Proliferating control cells were used as further control. Values were normalized to senescent control cells. **C)** Apoptosis levels measure by Annexin V incorporation in control and *sin-lncRNA* depleted IMR90 ER:RAS cells after 5 days of senescence induction. Proliferating cells were used as further control (n=3). **D)** RT-qPCR analysis of control and *sin-lncRNA* depleted TIG3-hTERT:BRAF cells after 3 days of senescence induction. Proliferating control cells were used as further control. Primer sets #3 (Suppl. Figure 2A) were used to amplify *sin-lncRNA*. Values were normalized to senescent control cells. **E)** BrdU staining and **F)** relative growth measured by crystal violet at 570nm absorbance of control and *sin-lncRNA* depleted TIG3-hTERT:BRAF cells after 5 days of senescence induction. Proliferating control cells were used as further control. Values were normalized to senescent control cells. **G)** Schematic representation of the CIRSPR/Cas9 strategy applied to remove *sin-lncRNA* genetic loci. sgRNAs guiding Cas9 protein are indicated (sgRNA_up and sgRNA_dw), as well as inner and outer primer couples to screen engineered cell clones (fwd/rev_in and fwd/rev_out, respectively) **H)** Gel images showing PCR products obtained using outer (left) or inner primer sets (right) to amplify the genome of TIG3 BRAF single cell clones after infection with the lentiCRISPR carrying *sin-lncRNA* sgRNAs and selection (clone#n). TIG3 BRAF fibroblasts infected with empty lentiviral vector (Empty) or infected with vector carrying *sin-lncRNA* sgRNAs but without single cell separation and puro selection (CRISPR_unsorted) were used as control.

**Suppl. Figure 4.**
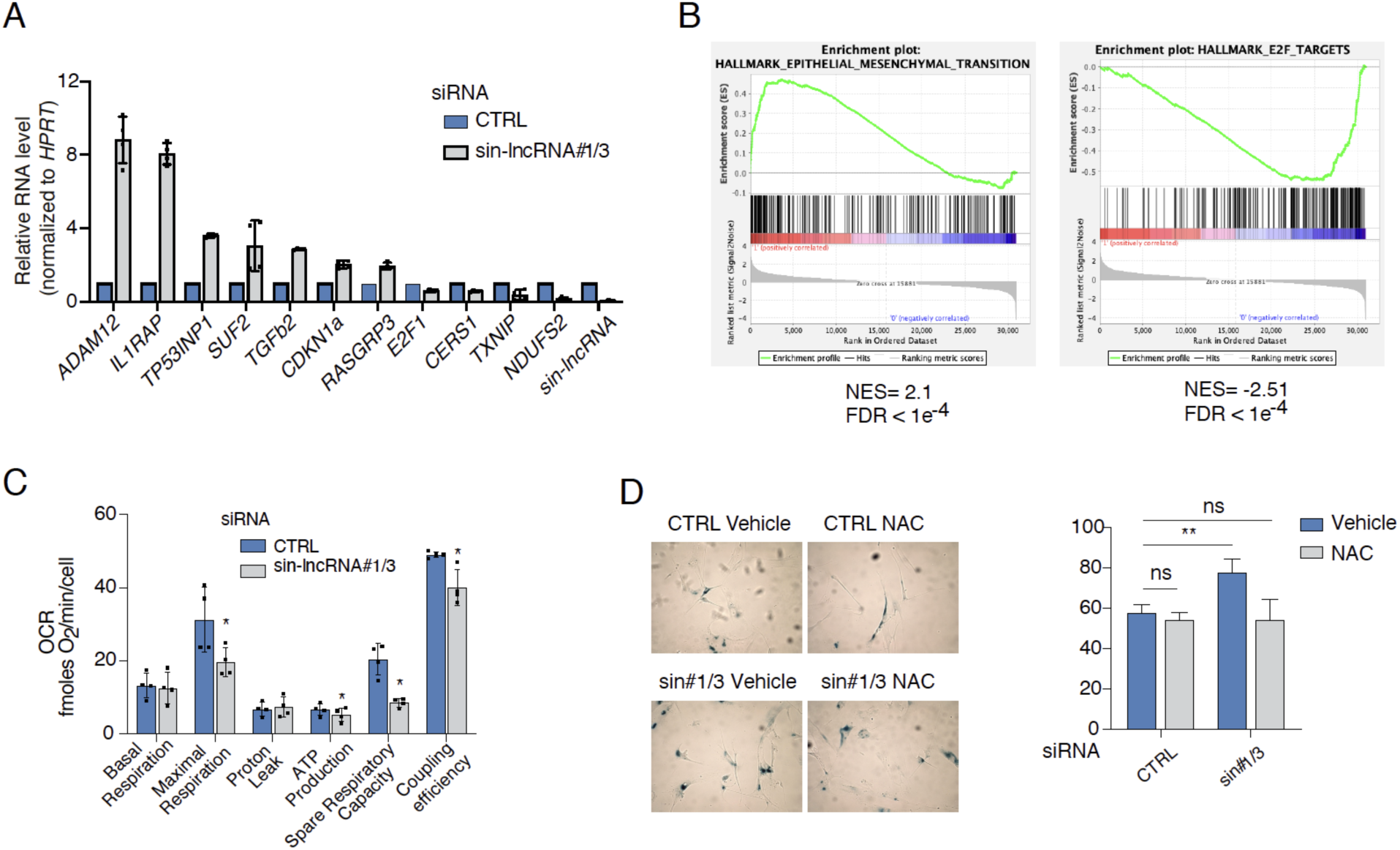
**A**) RT-qPCR analysis of the indicated transcripts in control and *sin-lncRNA* depleted IMR90 ER:RAS cells after 5 days of senescence induction. Proliferating control cells were used as further control. Values were normalized to senescent control cells. **B)** Gene Set Enrichment Analysis (GSEA) for genes belonging to epithelial to mesenchymal transition and E2F-targets pathways. **C)** Oxygen Consumption Rate (OCR) of control or *sin-lncRNA* depleted IMR90 ER:RAS senescent cells obtained using the Seahorse XFe96 Analyzer following injections of oligomycin, FCCP, and rotenone/antimycin A, as indicated in the Cell Mito Stress Test. Data is presented as mean ± s.d. (n = 4). Two tailed student’s *t*-test, *\*p<*0.05. **D)** Percentage of β-galactosidase positive cells in control or *sin-lncRNA* depleted IMR90 ER:RAS senescent cells treated with 5mM vehicle (DMSO) or N-acetyl-L-cisteine (NAC).

**Suppl. Figure 5.**
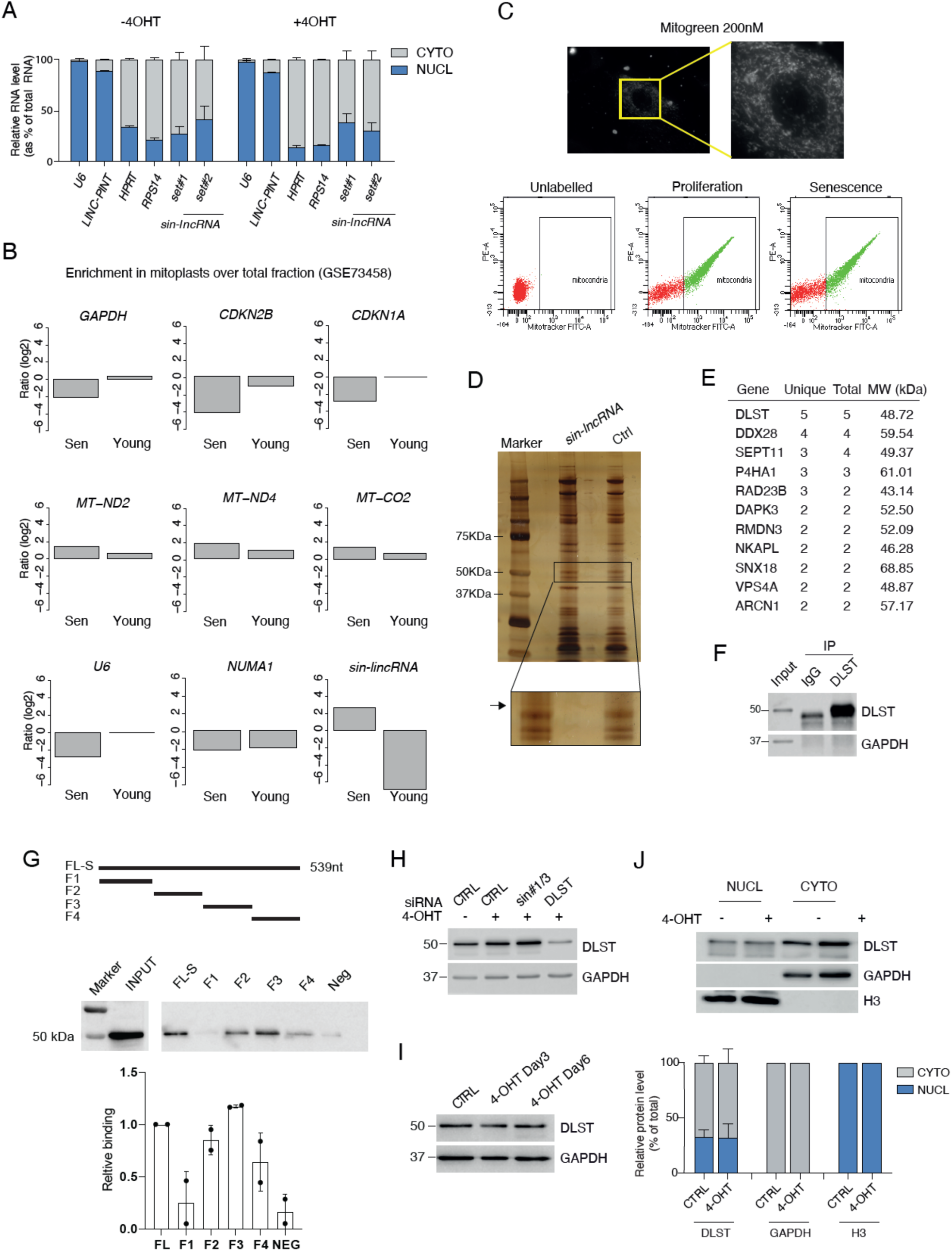
**A**) Percentage of nuclear and cytoplasmic RNA levels of *sin-lncRNA* measured by qRT–PCR after cell fractionation with two different set of primers (set#1 and set#2) in untreated or 4OHT-induced senescent cells. **B)** Enrichment of RNA in mitoplasts over the input (shown as log2) of the indicated genes in young and senescent WI38 fibroblasts analyzed from public available data (GSE73458). **C)** *Upper panel*, representative microscope images of mitochondrial staining in senescent cells using 200 nM of MitoTracker for 30min at 37°C. The white box shows the amplified area. Bottom panel, FACS plots indicating the isolation of mitochondria in proliferating and senescent cells. **D)** Representative image of the silver-stained gel. We show the band that was isolated and analyzed by mass spectrometry. **E)** List of top candidate proteins identified by mass spectrometry after pulldown with in vitro biotinylated *sin-lncRNA***. F)** Western blot analysis of the immunoprecipitated DLST using anti-DLST antibody or IgG as a negative control. GAPDH was used as control. (n=2)**. G)** Schematic representation of the fragments (full length (FL) and (F1-F5) used for the in vitro pulldown experiment. A nuclear lncRNA (CONCR) was used as negative control (Neg). *Bottom*, western blot to detect DLST using the fragments shown above. The graph shows the quantification of the binding relative to the full length (FL), (n=2). **H)** Western blot analysis of control IMR90 ER:RAS cells or cells transfected with siRNAs against *sin-lncRNA* or DLST. Proliferating control cells were used as further control. **I)** Western blot analysis of DLST protein levels in untreated IMR90 ER:RAS cells or treated for 3 or 6 days. GAPDH was used as a loading control. **J)** *Upper panel*, western blot analysis of DLST cellular fractionation in control and 4OHT-senescence indcued IMR90 ER:RAS. GAPDH and H3 were used as cytoplasmic and nuclear controls respectively. *Bottom panel*, percentage of nuclear and cytoplasmic distribution of DLST normalized to GAPDH (cyto) or H3 (nucl). The graph represents the mean ± s.d. (n=2).

**Suppl. Figure 6.**
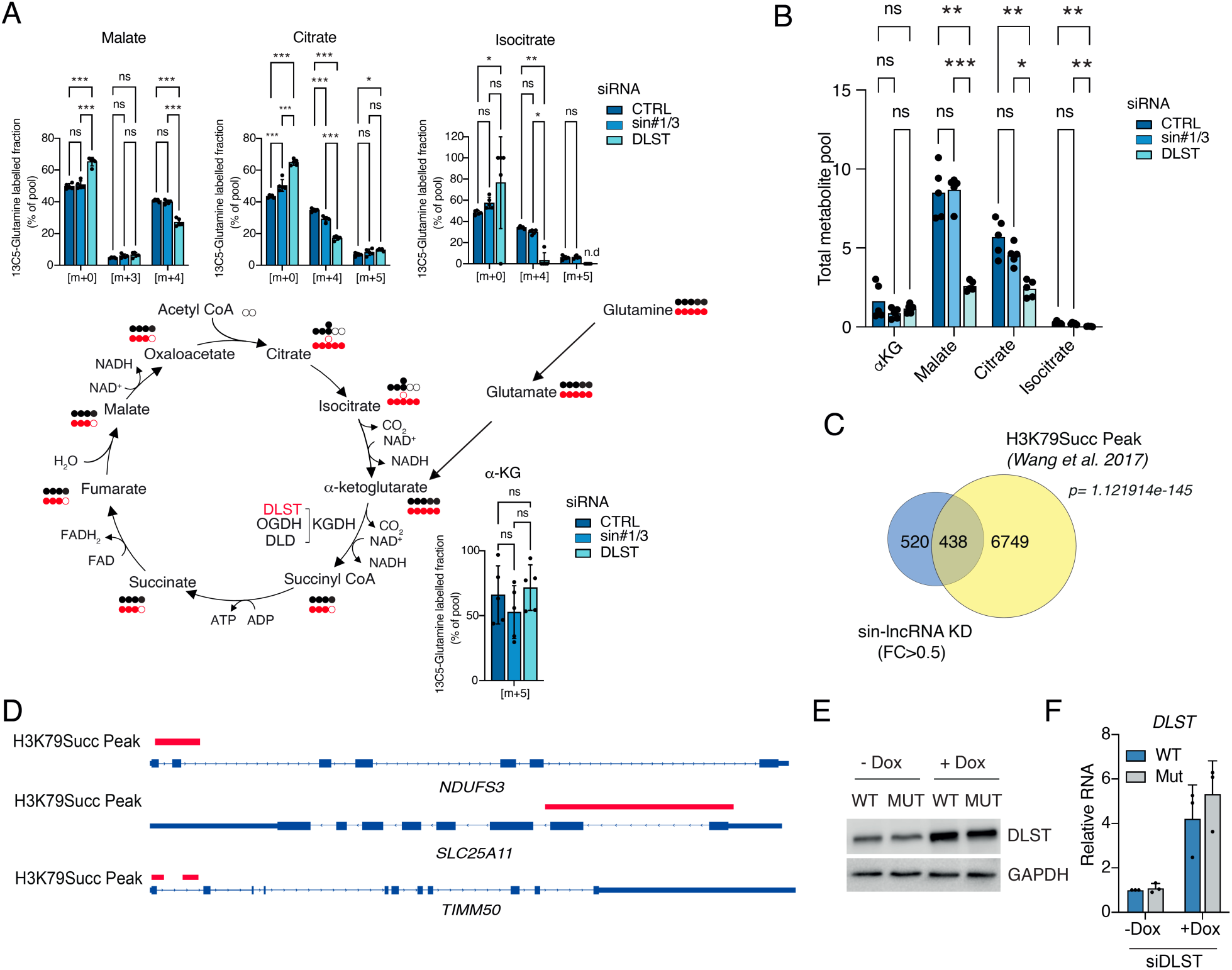
**A**) Carbon labelling TCA cycle intermediates from ^13^C_5_-Glutamine via oxidative (carbon marked with black) and reductive carboxylation (carbon marked with red) pathway. Labelling pattern of intracellular metabolites in senescent IMR90 ER:RAS fibroblasts transfected with siRNA control or siRNA against *sin-lncRNA* or DLST. Bar graphs represent the mean ± s.d. (n=5) for the isotopologues of most interest for each metabolite. Two tailed student’s *t*-test, *\*p<*0.05; **p<0.01; ***p<0.001. **B)** Bar plot showing differences in total metabolite pool size between control senescent cells or senescent cells depleted for *sin-lncRNA* or DLST (n=5). **C)** Venn diagram showing the overlap of genes deregulated in *sin-lncRNA* knock down cells (FC>0.5) and the H3K79Succ peaks. *p value* determined by hypergeometric test. **D)** IGV tracks showing the presence of H3K79Succ peak (red) in the corresponding genes. **E)** Western blot detection of DLST and GAPDH as housekeeping control in stable senescent IMR90 ER:RAS cells expressing a doxycycline-inducible wild type (WT) or NLS mutant (Mut) (R224A/K226E) DLST construct in the absence (-Dox) or presence (+Dox) of 100ng/ml of doxycycline for 72h (n=2). **F)** qRT-PCR of the *DLST* RNA level in the conditions described above (n=3).

**Suppl. Figure 7.**
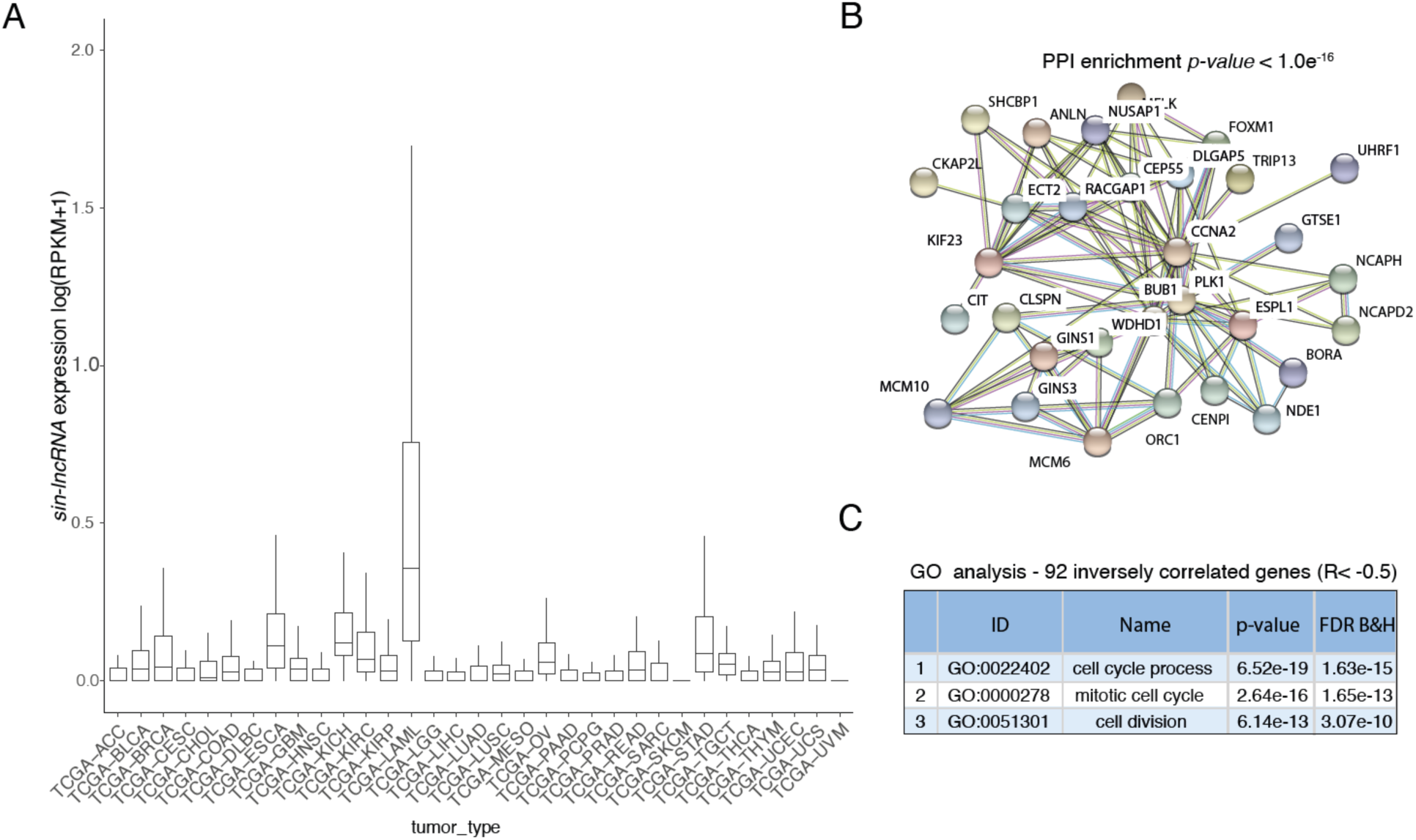
**A**) *Sin-lncRNA* expression (as FPKM + pseudocount value) across tumor samples from the TCGA database. **B)** Protein-Protein Interaction network (*p-value*<1.0e^-16^) by STRING analysis of genes inversely correlated with *sin-lncRNA* (92 genes; R< –0.5). **C)** Gene Ontology (GO) pathways associated with the inversely correlated genes.

## REFERENCES

1. Kuilman, T., Michaloglou, C., Mooi, W.J. & Peeper, D.S. The essence of senescence. Genes & development 24, 2463–2479 (2010).

2. Kuilman, T. et al. Oncogene-induced senescence relayed by an interleukin-dependent inflammatory network. Cell 133, 1019–1031 (2008).

3. Acosta, J.C. et al. Chemokine signaling via the CXCR2 receptor reinforces senescence. Cell 133, 1006–1018 (2008).

4. Coppe, J.P. et al. Senescence-associated secretory phenotypes reveal cell-nonautonomous functions of oncogenic RAS and the p53 tumor suppressor. PLoS biology 6, 2853–2868 (2008).

5. Coppe, J.P., Desprez, P.Y., Krtolica, A. & Campisi, J. The senescence-associated secretory phenotype: the dark side of tumor suppression. Annu Rev Pathol 5, 99–118 (2010).

6. Nardella, C., Clohessy, J.G., Alimonti, A. & Pandolfi, P.P. Pro-senescence therapy for cancer treatment. Nat Rev Cancer 11, 503–511 (2011).

7. Milanovic, M. et al. Senescence-associated reprogramming promotes cancer stemness. Nature 553, 96–100 (2018).

8. Dorr, J.R. et al. Synthetic lethal metabolic targeting of cellular senescence in cancer therapy. Nature 501, 421–425 (2013).

9. Gonzalez-Gualda, E., Baker, A.G., Fruk, L. & Munoz-Espin, D. A guide to assessing cellular senescence in vitro and in vivo. FEBS J 288, 56–80 (2021).

10. Kwon, S.M., Hong, S.M., Lee, Y.K., Min, S. & Yoon, G. Metabolic features and regulation in cell senescence. BMB Rep 52, 5–12 (2019).

11. Lee, A.C. et al. Ras proteins induce senescence by altering the intracellular levels of reactive oxygen species. The Journal of biological chemistry 274, 7936–7940 (1999).

12. Moiseeva, O., Bourdeau, V., Roux, A., Deschenes-Simard, X. & Ferbeyre, G. Mitochondrial dysfunction contributes to oncogene-induced senescence. Mol Cell Biol 29, 4495–4507 (2009).

13. Mattick, J.S. et al. Long non-coding RNAs: definitions, functions, challenges and recommendations. Nat Rev Mol Cell Biol 24, 430–447 (2023).

14. Statello, L., Guo, C.J., Chen, L.L. & Huarte, M. Gene regulation by long non-coding RNAs and its biological functions. Nat Rev Mol Cell Biol 22, 96–118 (2021).

15. Qureshi, I.A. & Mehler, M.F. Emerging roles of non-coding RNAs in brain evolution, development, plasticity and disease. Nat Rev Neurosci 13, 528–541 (2012).

16. Huarte, M. The emerging role of lncRNAs in cancer. Nat Med 21, 1253–1261 (2015).

17. Abdelmohsen, K. et al. Senescence-associated lncRNAs: senescence-associated long noncoding RNAs. Aging cell 12, 890–900 (2013).

18. Montes, M. et al. The lncRNA MIR31HG regulates p16(INK4A) expression to modulate senescence. Nat Commun 6, 6967 (2015).

19. Montes, M. et al. The long non-coding RNA MIR31HG regulates the senescence associated secretory phenotype. Nat Commun 12, 2459 (2021).

20. Montes, M. & Lund, A.H. Emerging roles of lncRNAs in senescence. FEBS J (2016).

21. Puvvula, P.K. et al. Long noncoding RNA PANDA and scaffold-attachment-factor SAFA control senescence entry and exit. Nat Commun 5, 5323 (2014).

22. Grossi, E. et al. A lncRNA-SWI/SNF complex crosstalk controls transcriptional activation at specific promoter regions. Nat Commun 11, 936 (2020).

23. Zhu, Y. et al. The long noncoding RNA glycoLINC assembles a lower glycolytic metabolon to promote glycolysis. Mol Cell 82, 542–554 e546 (2022).

24. Wang, P., Xu, J., Wang, Y. & Cao, X. An interferon-independent lncRNA promotes viral replication by modulating cellular metabolism. Science 358, 1051–1055 (2017).

25. Gandhi, M. et al. The lncRNA lincNMR regulates nucleotide metabolism via a YBX1 – RRM2 axis in cancer. Nat Commun 11, 3214 (2020).

26. Ding, C. et al. A T(reg)-specific long noncoding RNA maintains immune-metabolic homeostasis in aging liver. Nat Aging 3, 813–828 (2023).

27. Castello, A., Hentze, M.W. & Preiss, T. Metabolic Enzymes Enjoying New Partnerships as RNA-Binding Proteins. Trends Endocrinol Metab 26, 746–757 (2015).

28. Herranz, N. et al. mTOR regulates MAPKAPK2 translation to control the senescence-associated secretory phenotype. Nat Cell Biol 17, 1205–1217 (2015).

29. Vater, C.A., Bartle, L.M., Dionne, C.A., Littlewood, T.D. & Goldmacher, V.S. Induction of apoptosis by tamoxifen-activation of a p53-estrogen receptor fusion protein expressed in E1A and T24 H-ras transformed p53-/-mouse embryo fibroblasts. Oncogene 13, 739–748 (1996).

30 ENCODE Project Consortium. An integrated encyclopedia of DNA elements in the human genome. Nature, 489(7414):57–74 (2012)

31 Hoare, M. et al. NOTCH1 mediates a switch between two distinct secretomes during senescence. Nat Cell Biol 18, 979–992 (2016).

32. Innes, A.J. & Gil, J. IMR90 ER:RAS: A Cell Model of Oncogene-Induced Senescence. Methods Mol Biol 1896, 83–92 (2019).

33. Woods, D. et al. Raf-induced proliferation or cell cycle arrest is determined by the level of Raf activity with arrest mediated by p21Cip1. Mol Cell Biol 17, 5598–5611 (1997).

34. Nuskova, H. et al. Mitochondrial ATP synthasome: Expression and structural interaction of its components. Biochem Biophys Res Commun 464, 787–793 (2015).

35. Hutter, E. et al. Senescence-associated changes in respiration and oxidative phosphorylation in primary human fibroblasts. Biochem J 380, 919–928 (2004).

36. Zhang, J. & Zhang, Q. Using Seahorse Machine to Measure OCR and ECAR in Cancer Cells. Methods Mol Biol 1928, 353–363 (2019).

37. Martinez-Reyes, I. & Cuezva, J.M. The H(+)-ATP synthase: a gate to ROS-mediated cell death or cell survival. Biochim Biophys Acta 1837, 1099–1112 (2014).

38. Santamaria, G., Martinez-Diez, M., Fabregat, I. & Cuezva, J.M. Efficient execution of cell death in non-glycolytic cells requires the generation of ROS controlled by the activity of mitochondrial H+-ATP synthase. Carcinogenesis 27, 925–935 (2006).

39. Ni, R. et al. Mitochondrial Calpain-1 Disrupts ATP Synthase and Induces Superoxide Generation in Type 1 Diabetic Hearts: A Novel Mechanism Contributing to Diabetic Cardiomyopathy. Diabetes 65, 255–268 (2016).

40. Johnson, K.M. et al. Identification and validation of the mitochondrial F1F0-ATPase as the molecular target of the immunomodulatory benzodiazepine Bz-423. Chem Biol 12, 485–496 (2005).

41. Vizioli, M.G. et al. Mitochondria-to-nucleus retrograde signaling drives formation of cytoplasmic chromatin and inflammation in senescence. Genes & development 34, 428–445 (2020).

42. Velarde, M.C., Flynn, J.M., Day, N.U., Melov, S. & Campisi, J. Mitochondrial oxidative stress caused by Sod2 deficiency promotes cellular senescence and aging phenotypes in the skin. Aging (Albany NY) 4, 3–12 (2012).

43. Macip, S. et al. Inhibition of p21-mediated ROS accumulation can rescue p21-induced senescence. EMBO J 21, 2180–2188 (2002).

44. Staal, F.J., Roederer, M., Herzenberg, L.A. & Herzenberg, L.A. Intracellular thiols regulate activation of nuclear factor kappa B and transcription of human immunodeficiency virus. Proc Natl Acad Sci U S A 87, 9943–9947 (1990).

45. Leucci, E. et al. Melanoma addiction to the long non-coding RNA SAMMSON. Nature 531, 518–522 (2016).

46. Noh, J.H. et al. HuR and GRSF1 modulate the nuclear export and mitochondrial localization of the lncRNA RMRP. Genes & development 30, 1224–1239 (2016).

47. Tu, Y.T. & Barrientos, A. The Human Mitochondrial DEAD-Box Protein DDX28 Resides in RNA Granules and Functions in Mitoribosome Assembly. Cell Rep 10, 854–864 (2015).

48. Sheu, K.F. & Blass, J.P. The alpha-ketoglutarate dehydrogenase complex. Ann N Y Acad Sci 893, 61–78 (1999).

49. Street, L.A. et al. Large-scale map of RNA-binding protein interactomes across the mRNA life cycle. Mol Cell 84, 3790–3809.e8 (2024).

50. Shen, N. et al. DLST-dependence dictates metabolic heterogeneity in TCA-cycle usage among triple-negative breast cancer. Commun Biol 4, 1289 (2021).

51. Qattan, A.T., Radulovic, M., Crawford, M. & Godovac-Zimmermann, J. Spatial distribution of cellular function: the partitioning of proteins between mitochondria and the nucleus in MCF7 breast cancer cells. J Proteome Res 11, 6080–6101 (2012).

52. Wang, Y. et al. KAT2A coupled with the alpha-KGDH complex acts as a histone H3 succinyltransferase. Nature 552, 273–277 (2017).

53. Shen, N. et al. DLST-dependence dictates metabolic heterogeneity in TCA-cycle usage among triple-negative breast cancer. Commun Biol 4, 1289 (2021).

54. Bodineau C, et al. Two parallel pathways connect glutamine metabolism and mTORC1 activity to regulate glutamoptosis. Nat Commun. 2021 Aug 10;12(1):4814.

55. Frigola, J., Remus, D., Mehanna, A. & Diffley, J. F. ATPase-dependent quality control of DNA replication origin licensing. Nature 495, 339–343 (2013).

56. Alvarez-Abril, B., Garcia-Martinez, E. & Galluzzi, L. Platinum-based chemotherapy inflames the ovarian carcinoma microenvironment through cellular senescence. Oncoimmunology 11, 2052411 (2022).

57. Murphy, M.P. How mitochondria produce reactive oxygen species. Biochem J 417, 1–13 (2009).

58. Hamanaka, R.B. & Chandel, N.S. Mitochondrial reactive oxygen species regulate cellular signaling and dictate biological outcomes. Trends Biochem Sci 35, 505–513 (2010).

59. Davalli, P., Mitic, T., Caporali, A., Lauriola, A. & D’Arca, D. ROS, Cell Senescence, and Novel Molecular Mechanisms in Aging and Age-Related Diseases. Oxid Med Cell Longev 2016, 3565127 (2016).

60. Wei, Z. et al. CUL4B impedes stress-induced cellular senescence by dampening a p53-reactive oxygen species positive feedback loop. Free Radic Biol Med 79, 1–13 (2015).

61. Martinez-Reyes, I. & Chandel, N.S. Mitochondrial TCA cycle metabolites control physiology and disease. Nat Commun 11, 102 (2020).

62. Hansen, G.E. & Gibson, G.E. The alpha-Ketoglutarate Dehydrogenase Complex as a Hub of Plasticity in Neurodegeneration and Regeneration. Int J Mol Sci 23 (2022).

63. Tretter, L. & Adam-Vizi, V. Inhibition of Krebs cycle enzymes by hydrogen peroxide: A key role of [alpha]-ketoglutarate dehydrogenase in limiting NADH production under oxidative stress. J Neurosci 20, 8972–8979 (2000).

64. Hoogstraten, C.A. et al. Metabolic impact of genetic and chemical ADP/ATP carrier inhibition in renal proximal tubule epithelial cells. Arch Toxicol 97, 1927–1941 (2023).

65. Sutendra, G. et al. A nuclear pyruvate dehydrogenase complex is important for the generation of acetyl-CoA and histone acetylation. Cell 158, 84–97 (2014).

66. Kafkia, E. et al. Operation of a TCA cycle subnetwork in the mammalian nucleus. Sci Adv 8, eabq5206 (2022).

67. Li, W. et al. Nuclear localization of mitochondrial TCA cycle enzymes modulates pluripotency via histone acetylation. Nat Commun 13, 7414 (2022).

68. Chen, J. et al. Compartmentalized activities of the pyruvate dehydrogenase complex sustain lipogenesis in prostate cancer. Nat Genet 50, 219–228 (2018).

69. Mellid, S. et al. DLST mutations in pheochromocytoma and paraganglioma cause proteome hyposuccinylation and metabolic remodeling. Cancer Commun (Lond) (2023).

70. Castello, A. et al. Comprehensive Identification of RNA-Binding Proteins by RNA Interactome Capture. Methods Mol Biol 1358, 131–139 (2016).

71. Chatterjee, A. et al. RNA promotes mitochondrial import of F1-ATP synthase subunit alpha (ATP5A1) bioRxiv 2024.08.19.608659; doi: 10.1101/2024.08.19.608659

72. Ewald, J.A., Desotelle, J.A., Wilding, G. & Jarrard, D.F. Therapy-induced senescence in cancer. J Natl Cancer Inst 102, 1536–1546 (2010).

73. Pardella, E. et al. Therapy-Induced Stromal Senescence Promoting Aggressiveness of Prostate and Ovarian Cancer. Cells 11 (2022).

74. Pranzini, E., Pardella, E., Paoli, P., Fendt, S.M. & Taddei, M.L. Metabolic Reprogramming in Anticancer Drug Resistance: A Focus on Amino Acids. Trends Cancer 7, 682–699 (2021).

75. Tan, Y. et al. Metabolic reprogramming from glycolysis to fatty acid uptake and beta-oxidation in platinum-resistant cancer cells. Nat Commun 13, 4554 (2022).

76. Jiang, P., Du, W., Mancuso, A., Wellen, K.E. & Yang, X. Reciprocal regulation of p53 and malic enzymes modulates metabolism and senescence. Nature 493, 689–693 (2013).

77. Zhang, H. et al. LncRNA NEAT1 controls the lineage fates of BMSCs during skeletal aging by impairing mitochondrial function and pluripotency maintenance. Cell Death Differ 29, 351–365 (2022).

78. Vendramin, R., Marine, J.C. & Leucci, E. Non-coding RNAs: the dark side of nuclear-mitochondrial communication. EMBO J 36, 1123–1133 (2017).

79. Dieterle, F., Ross, A., Schlotterbeck, G. & Senn, H. Probabilistic quotient normalization as robust method to account for dilution of complex biological mixtures. Application in 1H NMR metabonomics. Anal Chem 78, 4281–4290 (2006).

80. Veselkov, K.A. et al. Optimized preprocessing of ultra-performance liquid chromatography/mass spectrometry urinary metabolic profiles for improved information recovery. Anal Chem 83, 5864–5872 (2011).

81. Marin-Bejar, O. et al. The human lncRNA LINC-PINT inhibits tumor cell invasion through a highly conserved sequence element. Genome Biol 18, 202 (2017).

82. Liao, Y., Smyth, G.K. & Shi, W. featureCounts: an efficient general purpose program for assigning sequence reads to genomic features. Bioinformatics 30, 923–930 (2014).

83. Langmead, B. & Salzberg, S.L. Fast gapped-read alignment with Bowtie 2. Nat Methods 9, 357–359 (2012).

84. Liberzon, A. et al. The Molecular Signatures Database (MSigDB) hallmark gene set collection. Cell Syst 1, 417–425 (2015).

85. Edgar, R., Domrachev, M. & Lash, A.E. Gene Expression Omnibus: NCBI gene expression and hybridization array data repository. Nucleic Acids Res 30, 207–210 (2002).

86. Perez-Riverol, Y. et al. The PRIDE database resources in 2022: a hub for mass spectrometry-based proteomics evidences. Nucleic Acids Res 50, D543–D552 (2022).

